# European Beech Spring Phenological Phase Prediction with UAV-derived Multispectral Indices and Machine Learning Regression

**DOI:** 10.1101/2022.12.30.522283

**Authors:** Stuart Krause, Tanja G.M. Sanders

## Abstract

The acquisition of phenological events play an integral part in investigating the effects of climate change on forest dynamics and assessing the potential risk involved with the early onset of young leaves. Large scale mapping of forest phenological timing using earth observation data, could facilitate a better understanding of phenological processes due to an added spatial component. The translation of traditional phenological ground observation data into reliable ground truthing for the purpose of the training and validation of Earth Observation (EO) mapping applications is a challenge. In this study, we explored the possibility of predicting high resolution phenological phase data for European beech (*Fagus sylvatica*) with the use of Unmanned Aerial Vehicle (UAV)-based multispectral indices and machine learning. Using a comprehensive feature selection process, we were able to identify the most effective sensors, vegetations indices, training data partitions, and machine learning models for phenological phase prediction. The best performing model that generalised well over various sites was the model utilising the Green Chromatic Coordinate (GCC) and Generalized Addictive Model (GAM) boosting. The GCC training data was derived from the radiometrically calibrated visual bands from a multispectral sensor and predicted using uncalibrated RGB sensor data. The final GCC/GAM boosting model was capable in predicting phenological phases on unseen datasets within a RMSE threshold of 0.5. This research shows the potential of the interoperability among common UAV-mounted sensors in particular the utility of readily available low cost RGB sensors. Considerable limitations were however discovered with indices implementing the near-infrared (NIR) band due to oversaturation. Future work involves adapting models to facilitate the ICP Forests phenological flushing stages.

## 1 Introduction

The concept of gathering data on the timing of leaf opening, flowering, fruiting and leaf fall alongside climatological observations “so as to show how areas differ” was first proposed by the Swedish botanist in his work *Philosphia Botinica* in 1751 (Linnaeus, 1751) and is still relevant today (Lieth, 1974). Phenological observations have historically assisted agriculture by means of predicting the timing for cultivation practices (Zhang, 2012) and in the last 100 years emerged as a scientific discipline (Schwartz, 2013). In recent times, phenological events are recognized as a bio-indicator for climate change (Menzel, 2002). Today, phenological data is a sensitive proxy for climate change investigation (Schwartz, 2013) due to an observed relationship between phenological timing and a changing climate. In particular, spring phenology is an indicator of climate change where observations are mirrored in temperature change (Menzel et al., 2006). Additionally, determining the heterogeneity of the phenological timings among individual phenotypes at the stand level can give insight in advantageous early spring flushing and the threat of late frost damage.

The Intergovernmental Panel on Climate Change (IPCC) has reported a 1.53°C increase in average land temperature for the period 2006 - 2015 in comparison to the 1850 - 1900 period (IPCC, 2018). Warmer temperatures, alongside changing precipitation patterns, have altered the growing seasons causing increased tree mortality (IPCC, 2018) however may also lead to increased carbon storage due to longer growing seasons (M. A. White et al., 1999). Linderholm (2006) reported that within the last few decades, the growing season has increased approximately 10 - 20 days and increasing average global temperatures (1.4 - 5.8 °C) in the next century could benefit some species and cause others to disappear.

A potential threat for European beech (*Fagus sylvatica L.*) in central Europe, when growing seasons are extended, is the earlier onset of budbreak and leaf-unfolding. When followed by sudden freezing temperatures, also known as episodic frost or “late frost”, newly unfolded leaves can be susceptible to the damage of cell tissue (Rubio-Cuadrado et al., 2021). Damage to the entire leaf or also partial damage, can have considerable effects on growth (Menzel et al., 2015; Sakai & Larcher, 1987). Although frost damage can potentially to some extent be compensated by the triggering of dormant or adventitious buds (Awaya et al., 2009), late frost damage can in effect shorten the growth season (Augspurger, 2009). Recently, in the spring of 2020, late frost damage was reported in the German state of Saxony (ca. 2.000 ha), in notably younger beech and oak stands without a protective upper canopy layer (Sachsenforst, 2020). For this reason, among others, provides the grounds for phenological modelling on a regional spatial scale which could prove beneficial in determining high risk areas where forest management practices can be adapted.

Temporal phenological models are typically determined by the timing of phenological occurrences (Zhao et al., 2013) and recorded in terms of day of year (DOY). For example, budbreak that occurred on May28th, 2020 would have the DOY of 149. Additionally, this day would have a temperature sum which is practically calculated as the sum of mean daily temperature, however better estimated in terms of the species-specific temperature-dependent modelling of “chilling days” and “thermal time” (officially known as “chilling unit” and “forcing unit” respectively) which is covered in detail by Menzel (1997) . Such models, are however susceptible to inaccuracies, as phenophases are not only affected by spring warming and winter chilling, but also photoperiod in terms of daily illumination and precipitation (Brügger & Vasella, 2018). The importance of accurate phenological observations and derived models could benefit as an early warning system for forest ecosystems that are under natural and anthropogenic stress (Brügger & Vasella, 2018). Additionally, accurate models can be used as predictors to simulate past, present and future phenological processes (Cleland et al., 2007). A better understanding of spatial and temporal variations of forest phenology on a regional level can give insights into the magnitude of climate change in terms of changing weather patterns and fluctuations of potential carbon fluxes (Cleland et al., 2007).

Visual phenological observations carried out by experts are conducted via long-term ground observations schemes (Menzel et al., 2006; Raspe et al., 2020; Vilhar et al., 2013) and enable data collection on stand and individual tree level. During such observations, data on for example crown condition is often recorded simultaneously and the root cause of potential tree damage can also be assessed. Such visual observations, although based on human subjectivity, are considered highly accurate, yet accuracy is limited to the experience of the observer and results could vary when multiple observers are involved. Visual ground observations are also labor- and time-intensive (Menzel 2015), restricted in spatial coverage, yet a valuable source of ground truthing. Standardized phenological observations carried out at Level II intensive monitoring plots, in accordance with ICP Forests (http://icp-forests.net) are comprised of over 600 plots in selected forest ecosystems throughout Europe and provide valuable information regarding the actual condition of individual trees (Raspe et al., 2020).

Terrestrial observation methods based on webcams and automated cameras mounted on towers have become increasingly popular for monitoring phenology in forest ecosystems (Ahrends et al., 2008; Alberton et al., 2017; Menzel et al., 2015; Sonnentag et al., 2012) and have developed into standardized networks such as the *PhenoCam* network (Brown et al., 2016; S. T. Klosterman et al., 2014) . Typically, daily and even hourly images are recorded throughout the growing season providing quantitative multispectral data that can deliver the exact timing of the phenophases of a particular species. As with the traditional ground observations, digital repeat photography observations are also limited in spatial coverage however play an important role in the validation of satellite observation platforms (N. Li et al., 2020) which assist with the mapping and modelling of phenological metrics on a global scale (Zeng et al., 2020).

Since the first *Phenology Satellite Experiment* in the early 1970’s (Dethier et al., 1972), satellite-based remote sensing has become an important tool for the study of phenology at multiple spatial scales (Kowalski et al., 2020). Current open access satellite remote sensing platforms such as Landsat 8 and Sentinel 2a/b deliver imagery with global coverage at a temporal and spatial resolution of 16 days and 30 meters (Landsat, 2022) and 5 days and 10 meters (Copernicus, 2022) respectively. The MODIS Global Land Cover Dynamics Product (Collection 5) delivers land surface phenology (LSP) information on a global scale at a 500 m spatial resolution based on 8-day input data (Friedl et al., 2010; Ganguly et al., 2010). Despite the high resolution capabilities of open access satellite data, there still lies the challenge of linking plot level measurements to satellite derived pixel values (White et al., 2014) as an occurrence of phenological heterogeneity among individuals within a single observation plot can take place (S. Klosterman et al., 2018) among other reasons.

The use of Unmanned Aerial Vehicles (UAVs) for the purpose of enhancing phenological observations has grown in recent years and is proving a promising method in filling the gap between terrestrial- and satellite-based phenological observation systems. The challenge lies in that a typical terrestrial observation plot will cover a limited amount of Satellite pixels. Berra (2019) demonstrated that phenological events within a single Landsat pixel can vary significantly (R^2^ < 0.50) in comparison to that of UAV-derived phenometrics. Additionally, Satellite pixels will not necessarily be capable of taking in account for microclimatic variability (S. Klosterman et al., 2018) as would be the case for plots where individual tree phenology is inhomogeneous. This brings forth the notion that UAV data trained from localized observation plots could scale localized observations to cover more area, thus enabling the training of more satellite pixels. Along these lines, Atkins et al. (2020) showed that UAV and terrestrial camera systems can be used complementary to obtain high resolution phenological data.

The conversion of acquired UAV data into phenological metrics has however it’s challenges with regards to sensor calibration during various environmental conditions as well as the magnitude of the processing workloads involved in the preparation of analysis-ready yearly multi-temporal high-resolution datasets. The aim of flight campaigns has two prominent purposes: the first being to determine the onset of spring leafing out (“flushing”) as is carried out at intensive forest monitoring (Level II) plots or more intensive observations such as at the Britz Reseach Station (north-eastern Germany) where each individual phase is observed; the second purpose being for the acquisition of training data which is representative of the complete phenophase range. As opposed to flushing observations, the phenophases, described in further detail in section 3, are more challenging to acquire as the individual phase occurrences may not align with fair weather windows conducive to flight missions and would involve more flights spanning over a month of required field work. A typical spring phenology campaign could be comprised from 6 to even 15 flight missions for the spring green-up and autumn senescence respectively depending on the phenophase resolution required. Additionally, such campaigns would also involve repeated missions at various observation plots in order to acquire representative data of a particular region and tree species. For more intensive monitoring programs, a repeat time of at least three days or even less during the early phenophases would be required to account for the timing of “bud burst”. Such data acquisition methods do however require automatic methods for the organization and preparation of data and the use of “explainable” machine learning (ML) algorithms (Belle & Papantonis, 2021) alongside parallel processing is an imperative if the acquired data mass is to be translated into useful and accurate phenological information. Automated methods however, should be implemented cautiously as the complexities of the phenological development of an individual tree are best assessed by experts using standardised qualitative methods which would prove useful for the production of expert-based annotations of single-shot aerial images as training for future large-scale data-driven deep learning applications. The application of data-hungry deep learning algorithms for the purpose of forest remote sensing (Diez et al., 2021; Kattenborn et al., 2021) in particular for phenological observations poses the problem of limited data and also the fact that the “view from above” is simply different from that from below the crown level causing uncertainty between the linkage of ground truthing to earth observation (EO) platforms. An automated operational deep learning infrastructure for forest monitoring is a hopeful goal for the future, but until then the implementation of “explainable” traditional ML algorithms as explored in this study and are necessary in avoiding the overfitting of models based on smaller datasets and can aid in developing a better understanding of forest processes as well as available features from the near-field aerial view provided by UAV-mounted sensors.

The use of ML algorithms for remote sensing applications, in particular image classification has become an ever-increasing topic of study in recent years. With regard to numerous publications, ML algorithms are proving to be effective at modelling complex datasets with higher accuracy than that of traditional parametric methods (Maxwell et al., 2018). An array of ML algorithms are available which can be implemented as classifiers or in regression mode for continuous data (Kuhn, 2008). Typical algorithms in usage in terms of supervised classifiers tend towards Support Vector Machines, Random Forests and Neural Networks (Lary et al., 2016; Schulz et al., 2018), however various ML techniques designed for specific applications exist.

With regards to the modelling of plant phenology, the use of remote sensing-based ML algorithms for the purpose of phenological modelling is somewhat limited. A particular interesting aspect of ML and associated feature engineering is the incorporation of features such as pixel values derived from vegetation indices combined as training data with meteorological data. Such implementations can provide a spatial aspect to the influence of temperature and precipitation on phenological processes. Czernecki et al. (2018) tested various regression-based ML techniques where the implementation of metrological indices alongside remote sensing products showed promising results for the estimation for spring and fall phenophases. In a study carried out by Dai et al. (2019), a gradient boosting decision tree (GBDT) model was used to assess the temperature sensitivity of long-term leaf unfolding data showing a potential underestimation of traditional phenological models. Further uses of UAV-based ML modelling was carried out by Park et al. (2019) applying stochastic gradient tree boosting for the monitoring individual tropical tree phenology by incorporating various RGB-based textures and vegetation indices.

In this study we explore the possibility to automate phenological phase extraction for european beech using UAV-based multispectral data and ML algorithms. At first, phenological trends originating from long-term field observation since 2006 will be explored at the Britz research station. To follow, the airborne multispectral data as well as derived indices from the years 2019-2021 will be analysed for a correlation analysis and polynomial fitting with field observations for the purpose of feature selection. Moreover, ML techniques in regression mode will be implemented to train the models based on training data from 2019 and 2020 and tested against the unseen spring phenological phases of 2021. Finally, the selected model is trained with variations of data subsets based on the year of origin and rigorously tested on fresh data from 2022 and data originating from older beech stands (> 50 years) and different regions.

## 2 Methods

### 2.1 Study Site

UAV and ground-based phenological observation for this study were carried out at the Britz intensive forest monitoring research station (shorturl.at/anoqT) located in the lowlands of north-eastern Brandenburg, Germany. Brandenburg lies in between oceanic and continental climate zones and belongs to the young moraine landscape of the *Weichsel* glacial period. The soil at the site is locally known as a “*Finowtaler Sandbraunerde*” (Schulze & Kopp, 1998) and rated as a *Dystric Cambisol* derived from *Pleistocene* sand deposits with a *moder* organic layer (Don et al., 2019). Average yearly temperature and precipitation for the region has been recorded at approximately 8.9 °C and 570 mm (Riek, 2004) respectively. Datasets acquired from other sites, for the purpose of testing Models with unseen data, were acquired from “Kahlenberg” near the Britz research station, and the “Black Forest” located north of Freiburg in south-western Germany.

The Britz research station was initially designed in 1972 for intensive forest hydrology research with 9 large-scale lysimeters planted with a mixture of deciduous and coniferous species (Müller 2010). The site is also equipped with an array of digital and analog dendrometers, and various other sensors for sapflow and soil moisture measurement as well as meteorological data. Recently, the research station as undergone a major digitalization overhaul where sensor data is automatically synced to the cloud and individual trees are mapped at a sub-decimeter accuracy.

### 2.2 Phenological Ground Observations

Ground-based phenological observations have been carried out at the Britz research station since 2006 over all nine plots as well as at various satellite plots. UAV-based phenological missions have been implemented at the research site since 2018 with the aid of on-board multispectral sensors. As of 2021, phenological observations are additionally carried out with tower-based phenology cameras.

In this study, the spring phenology of 14 trees of an approximately 50-year-old European Beech (*Fagus sylvatica*) stand was recorded via traditional observations methods alongside UAV-based missions with an offset of a minimum of ± 3 days. After 2020, traditional observations were carried out synchronous with UAV missions.

Traditional terrestrial phenological observations for beech recorded at the Britz research station are based on spring green-up phases (0-5) and foliation (%), and discoloration (%) and foliation (%) for fall senescence. The spring phases are comprised of 5 phases which are based on a combination of tree phenological observation techniques (Baumgartner, 1952; Brügger & Vasella, 2018; Forstreuter, 2002; Schüler, 2012; Schwartz, 2013). This method possesses a possible phase range between 0.0 and 5.0, where decimal values are used to interpret in between the phases. For example, in the case that 80 % of the observed tree crown has reached phase 4 then this will be recorded as phase 3.8. The “*Britzer*” method of spring phenological phase assessment is unique in the sense that it emphasizes the early onset of bud development with reference to the swelling of buds (phase 0.5) in early spring as well as the hardening and darkening of leaves in phase 5. Table 1 describes the various phases implemented with the “*Britzer*” method alongside other well-known techniques. The differentiation between phase 4.0 and 5.0 is implemented with the *Britzer* method and only considered with the Malisse/Schüler steps (Malaisse, 1964; Schüler, 2012). Figure 1 gives a photographic representation of the *Britzer* method phases.

**Figure 1:**
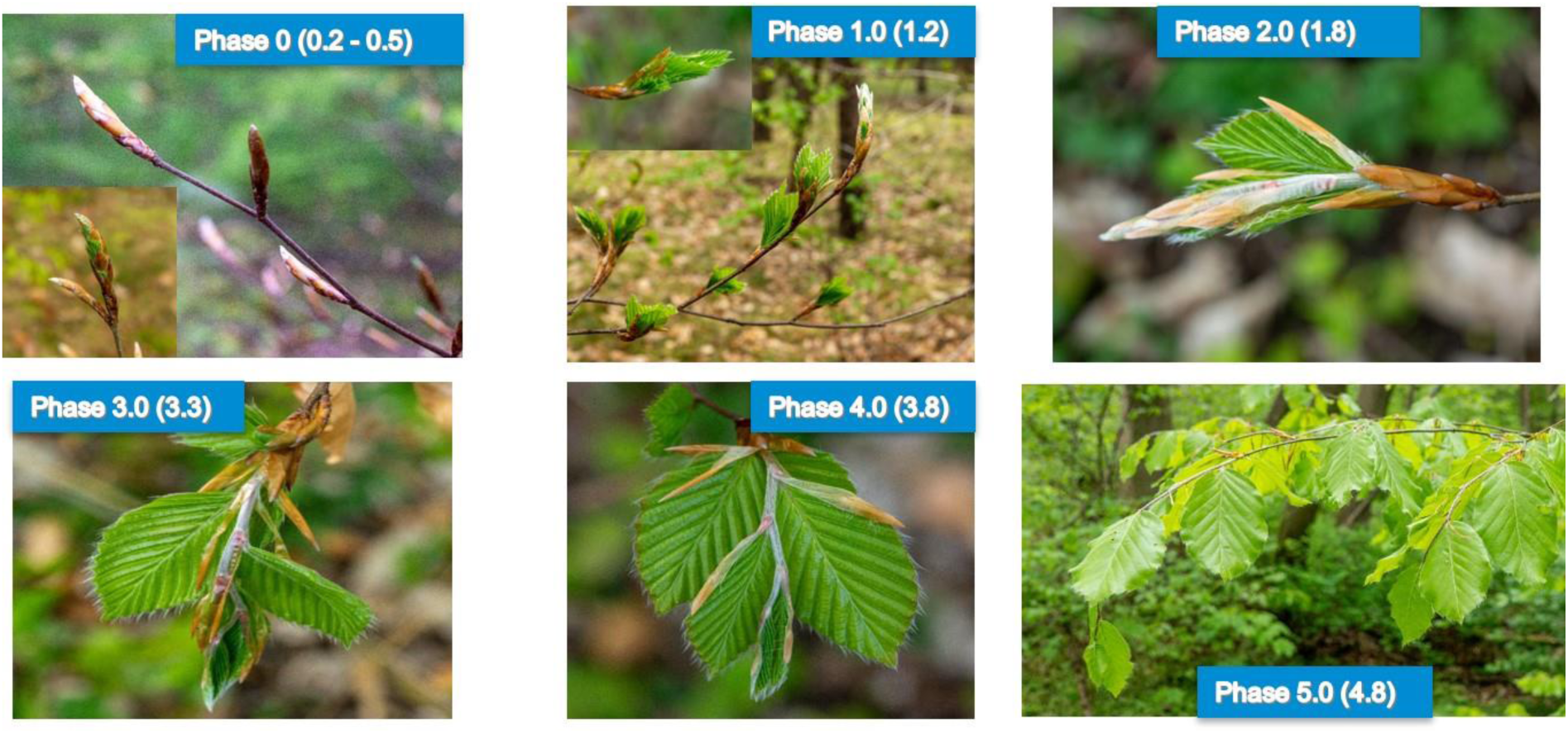
Photographic representation of the “Britzer” phenological phases.

**Table 1:** Overview of the various tree phenology observation methods for spring leafing out.

| **Forstreuter**<br>**(Stages)** | **Malaisse/Schüler**<br>**(Steps)** | **LFE**<br>**(Phases)** | **ICP Forests**<br>**(Flushing)** | **Britzer**<br>**(Phases)** | **Description** |
| --- | --- | --- | --- | --- | --- |
| 0A | 1 | 1 | 1 | 0 | Buds established from previous year |
| 0A | 1 | 1 | 1 | 0.2 | Buds in Winter dormancy |
| 0B | 2 | 2 | 1 | 0.5 | Swelling |
| 0C | 2 | 2 | 2 | 0.8 | First buds are bursting (ICP “flushing” = infrequent or slight) |
| 1D | 3 | 3 | 2-4 | 1.0 | Budburst (full tree) |
| 2E | 4 | 4 | 2-4 | (1.5) |  |
| 3 | 5 | 5 | 5 | 2.0 | Young wrinkled leaf visible |
| - | 6 | - | - | 3.0 | Young leaf less wrinkled and long shoot begins to lengthen. |
| - | 6 | - | - | 4.0 | Long shoot fully developed and pilose. Leaves still soft and pilose. |
| - | 7 | - | - | 5.0 | Long shoot almost glabrous and leaves hardened as well as dark green, less pilose |

Alongside the phenological phases, the *Britzer* method also takes foliation percentage into account (0-100 %) which is not directly harmonisable with the ICP Forests flushing method (Raspe et al., 2020). The ICP Forests method is shown in Table 1 for comparison purposes only and should not be taken as a direct method for conversion. Important to emphasis, is that the *Britzer* method records foliation as the coverage in % of fully developed leaves whereas the ICP Forests method of flushing records the coverage of green foliation and are not directly harmonisable.

### 2.3 UAV Multispectral Image Acquisition

The UAV remote sensing platform used for this study was comprised of two Unmanned Aerial Systems (UAS) customized for dual sensor RGB and multispectral image acquisition. The UAS implemented in 2019 was comprised of a *DJI Phantom 4 Professional Obsidian* (dji.com) with a RGB sensor (mechanical shutter) and a *Micasense Rededge-MX* (micasense.com) multispectral sensor mounted with a custom 3D-printed gimbal (droneparts.de). The UAS implemented after 2020 was a *DJI Martice 210 RTK* (dji.com) with built-in Real-Time-Kinematic capabilities which applies real-time corrections through a Network Transport of RTCM via Internet Protocol (NTRIP). Mounted on the *Matrice 210 RTK* was a *Zenmuse X7* RGB sensor alongside a *Micasense Altum* (micasense.com) multispectral sensor. Specific sensor details for the two multispectral and two RGB sensors can be found in Table 2 . Table 3 displays the wavelength and bandwidth information for both of the multispectral sensors. The Longwave Infrared (LWIR) band was not implemented for this study.

**Table 2:**
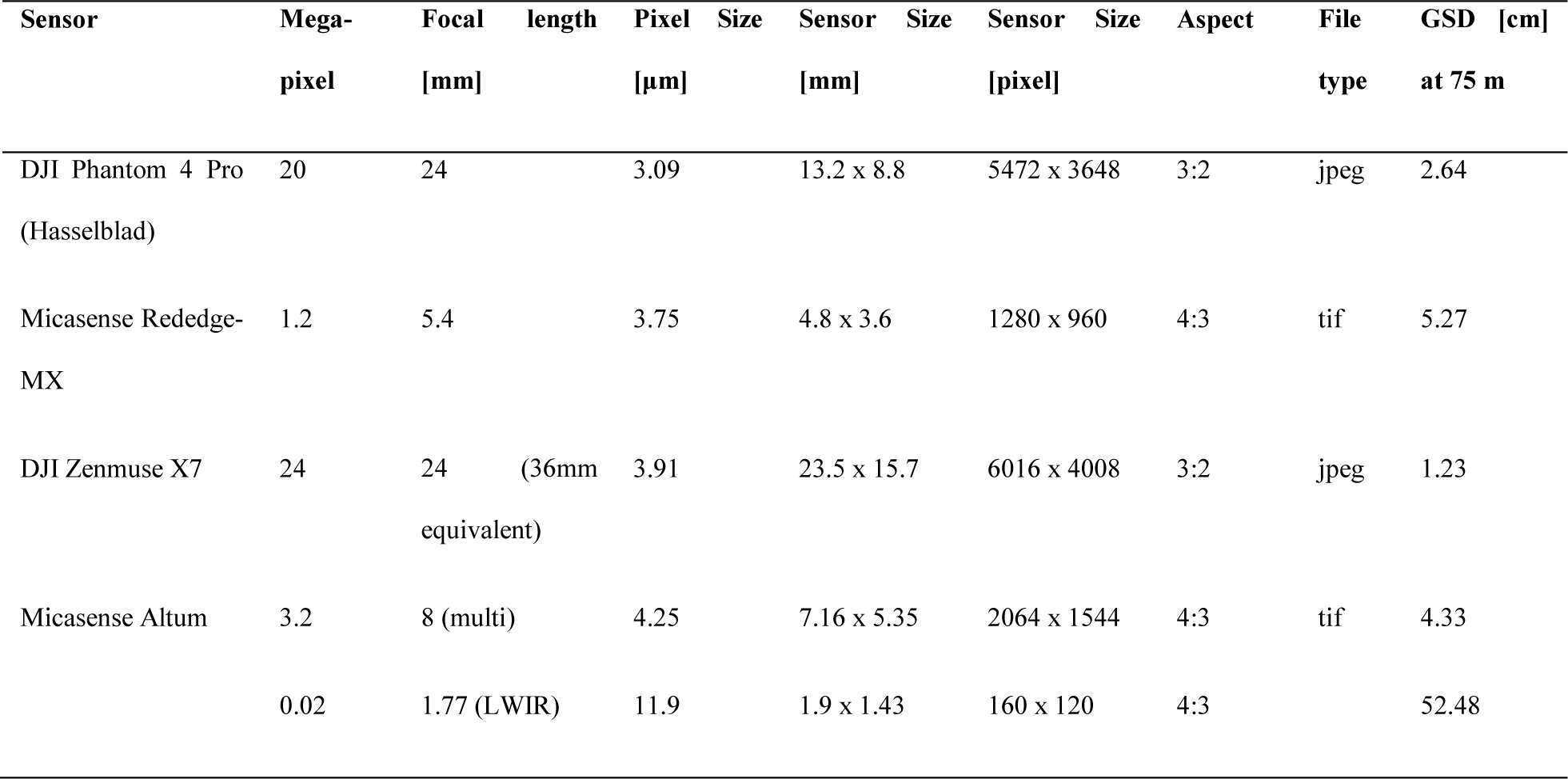
Overview of the sensor parameters used in this study.

| Sensor | Mega-<br>pixel | Focal<br>length<br>[mm] | Pixel<br>Size<br>[μm] | Sensor<br>Size<br>[mm] | Sensor<br>Size<br>[pixel] | Aspect | File<br>type | GSD [cm]<br>at 75 m |
| --- | --- | --- | --- | --- | --- | --- | --- | --- |
| DJI Phantom 4 Pro<br>(Hasselblad) | 20 | 24 | 3.09 | 13.2 x 8.8 | 5472 x 3648 | 3:2 | jpeg | 2.64 |
| Micasense Rededge-MX | 1.2 | 5.4 | 3.75 | 4.8 x 3.6 | 1280 x 960 | 4:3 | tif | 5.27 |
| DJI Zenmuse X7 | 24 | 24<br>(36mm<br>equivalent) | 3.91 | 23.5 x 15.7 | 6016 x 4008 | 3:2 | jpeg | 1.23 |
| Micasense Altum | 3.2 | 8 (multi) | 4.25 | 7.16 x 5.35 | 2064 x 1544 | 4:3 | tif | 4.33 |
|  | 0.02 | 1.77 (LWIR) | 11.9 | 1.9 x 1.43 | 160 x 120 | 4:3 |  | 52.48 |

**Table 3:** Wavelength and bandwidth for the Micasense Rededge-MX and Altum multispectral sensors.

| Band<br>Name | Micasense Altum |  | Micasense Rededge-MX |  |
| --- | --- | --- | --- | --- |
|  | Center |  | Center |  |
|  | Wavelength<br>(nm) | Bandwidth [nm] | Wavelength<br>(nm) | Bandwidth (nm) |
| Blue | 475 | 32 | 475 | 20 |
| Green | 560 | 27 | 560 | 20 |
| Red | 668 | 14 | 668 | 10 |
| Red Edge | 717 | 12 | 717 | 10 |
| Near IR | 842 | 57 | 840 | 40 |
| LWIR | 11 μm | 6 μm | - | - |

Phenology-based image acquisition missions at the Britz research station were carried out every spring (since 2019) typically synchronous with ground observations. A mission is carried shortly before budburst when buds begin swelling (0.5) and thereafter cycling in a maximum of 3 days during the fast-developing phases of 0.5 to 3. After the third phase, flights and ground observations are limited to bi-weekly acquisition days due to time constraints. Flight mission were carried out near solar noon (± 90 minutes) and calibration panel images (micasense.com, 2022) taken before and after missions as well as the acquisition of Downwelling Light Sensor (DLS) information for each individual multispectral image. After 2020, special care was taken to acquire calibration panel images during moments when the sun was the least affected by cloud cover, especially when the sun was revealed during the flight mission. In order to insure the capture of as many phases as possible, flight missions were carried out regardless of cloud cover, and refrained from during precipitation and for the most part winds over 12 km/h (3.3 ms).

Both *Micasense* multispectral sensors were set to capture images with an intervalometer set at 2 seconds and automatic exposure. The RGB sensors were typically set on shutter priority with a speed of 1/800^th^ of a second. Due to a slow trigger speed of the *Zenmuse X7*, flight mission speed was required to be reduced to 3 m/s to insure a forward overlap of at least 80 %.

### 2.4 Data Processing

After each flight campaign, images and metadata were stored on an external hard drive. Naming conventions were based upon a running ID, date, station/district, parcel number, and sensor. After the storage procedure, new similar folder names were created based on individual missions with only the selected images required for the Structure from Motion (SfM) software. These folders were then uploaded to an institute-based Linux High Performance Computing (HPC) cluster. Photogrammetric products were then produced in *Agisoft Metashape* (v1.7.5) with a semi-automated loop in *Python* where the script is interrupted to manually select Ground Control Points (GCPs) in the RGB and multispectral imagery (see Figure 2). The GCPs insured the repeated usage of crown segmentations throughout the time-series. The script loops through all folder names per Year which also implemented the naming conventions for each individual photogrammetric products such as the point cloud, Digital Surface Model (DSM), and orthomosaic.

**Figure 2:**
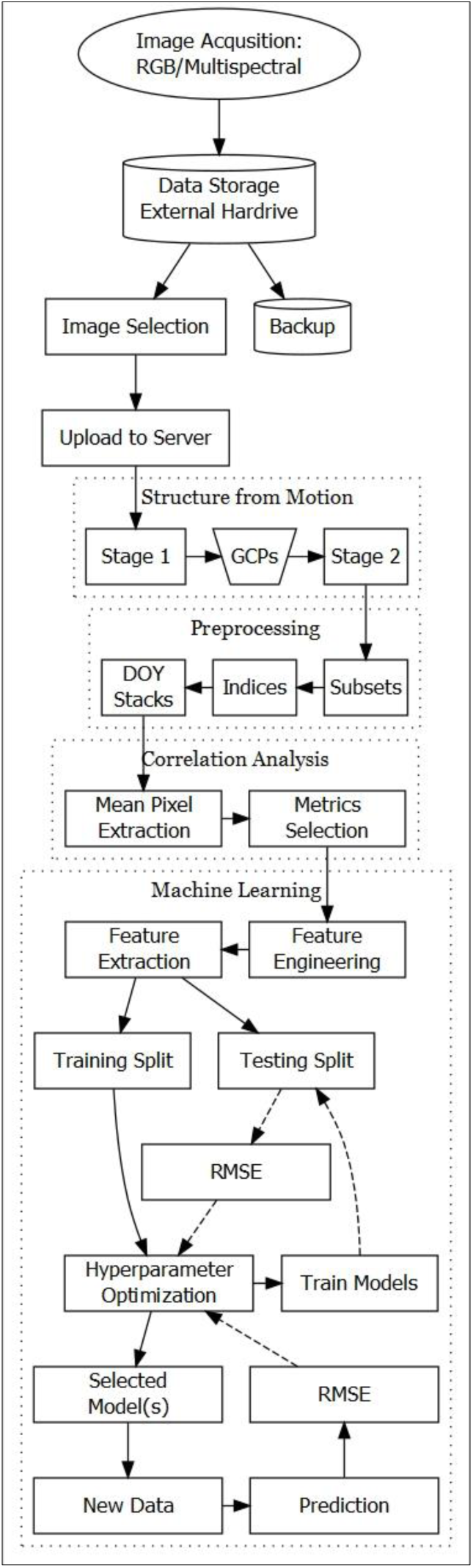
Workflow depicting the various steps from data acquisition to model evaluation.

Further processing was carried out in *R* (R Core Team, 2022) where spectral information was extracted from each individual phenology tree. Each dataset was subsetted to an Area-of-Interest (AOI) with a minimum boundary containing all of the trees used for the ground-based phenological observations. Preceding the calculation of the vegetation indices, the datasets were resampled to 0.01 m for data derived from the RGB sensors, and 0.03 m for the multispectral sensors. This enabled layer stacking based on the DOY and sensor. For models which implemented a fusion of both RGB and multispectral data, a resolution of 0.03 m was used. Shapefiles of the manually delineated tree crowns were implemented to extract the mean pixel values from all DOY layer stacks and stored in tabular format for further analysis. For the most part, the workflow was required to be automated due to copious amount of data (e.g. 14 tree crowns from 13 – 15 epochs per year from two sensors). Figure 2 shows the full workflow from image acquisition to model evaluation.

### 2.5 Vegetation Indices

Vegetation indices are widely used in remote sensing and can not only aid in detecting green vegetation traits but also reduce the effects of irradiance and variations in atmospheric transmission (Jones & Vaughan, 2010; Liang & Wang, 2020). Table 4 displays the indices used in this study. The Green Chromatic Coordinate (GCC) and the Normalized Green Red Difference Index (NGRDI) are the two indices which are derived from the visible part of the electromagnetic spectrum and available for typical consumer grade RGB sensors. Indices denoted with a “UC” (e.g. GCC_UC or GNRDI_UC) depict an index derived from bands that did not undergo radiometric calibration (uncalibrated).

**Table 4:** Vegetation Indices (VI) used in the study. NIR = Near-infrared.

| Name | VI | Formula | References |
| --- | --- | --- | --- |
| Green Chromatic Coordinate | GCC | $(\text{Green} / \text{Red} + \text{Green} + \text{Blue})$ | (Gillespie et al., 1987) |
| Normalized Green Red<br>Difference Index | NGRDI | $(\text{Green} - \text{Red}) / (\text{Green} + \text{Red})$ | (Tucker, 1979) |
| Normalized Difference<br>Vegetation Index | NDVI | $(\text{NIR} - \text{Red}) / (\text{NIR} + \text{Red})$ | (Rouse et al., 1974) |
| Normalized Difference Red-<br>edge Red Index | NDRE<br>(ReNDVI) | $(\text{Rededge}_{717} - \text{Red}_{668}) /$<br>$(\text{Rededge}_{717} + \text{Red}_{668})$ | Wang (2021) |
| Normalized Difference Red-<br>edge Index | NDREI | $(\text{NIR} - \text{Rededge}) / (\text{NIR} + \text{Rededge})$ | (A. Gitelson & Merzlyak, 1994b) |
| Green Normalized Difference<br>Vegetation Index | GNDVI | $(\text{NIR} - \text{Green}) / (\text{NIR} + \text{Green})$ | (A. A. Gitelson et al., 1996) |
| Enhanced Vegetation Index | EVI | $2.5 * (\text{NIR} - \text{Red}) / (\text{NIR} + 6 * \text{Red} - 7.5 * B + 1)$ | (Huete et al., 2002) |
| Normalized Difference Water<br>Index | NDWI | $(\text{Green} - \text{NIR}) / (\text{Green} + \text{NIR})$ | (McFeeters, 1996) |

The Normalized Difference Red-edge Index developed originally by Gitelson and M. Merzlyak (1994) was labeled NDREI in conformity with Hunt et al. (2013). The NDREI is sometimes labelled NDRE (Barnes et al., 2000), however in this study for practical reasons we use the abbreviation NDRE to depict the use of the Red-edge and Red bands rather than the Near-infrared and Red-edge bands. Another option could be to use the abbreviation ReNDVI used by Wang (2021).

The air temperature feature (AIRTEMP) was created from a summation of the daily air temperature above 0 °C (“warming days”) since January 1^st^ of a given year and implemented in this study for experimental purposes.

### 2.6 Feature Selection

With the aim of selecting appropriate features for the modelling process, a correlation analysis among independent and dependent variables was implemented. The correlation coefficient is scaleless and represented with the letter *r* which is interpreted with values between −1 and 1 where −1 would represent a perfect negative correlation in that the two variables have an inverse relationship and 1 represents a perfect linear relationship. 0 would depict the situation where both variables have no linear relationship (McClave & Sincich, 2018). In this study, the non-parametric Spearman correlation matrix was implemented for each of the indices in relation to the phenological phase and foliation. A test for multicollinearity was carried out which explored the between-variable correlation among predicting features (Kuhn & Johnson, 2019) for the purpose of improving feature selection for multivariate models. The use of polynomial regression models of the first to fifth order were also used to further evaluate individual features during the selection process.

### 2.7 Statistical Analysis and Machine Learning

The ML regression models implemented in this study were that of Generalized Additive Models (GAMs), Boosted GAMs and Gradient Boosting Machine (GBM). Model training was carried out in *R* with the *caret* package (Kuhn et al., 2022). Models were trained with a 80/20 training/validation split, scaling, and 10-Fold cross validation. The ML modelling process was divided into three main stages: 1) training/validation split of the subset variations of the 2019/2020/2021 datasets and test for correlation and polynomial fitting for feature selection purposes 2) Models trained with the full combined 2019/2020 datasets and tested with the 2021 dataset 3) further testing with selected models trained with subset variations of the 2019/2020/2021 datasets and tested against new single epoch datasets acquired from unseen data. The concept behind the subset variations is for the purpose of identifying which years are propagating error.

ML Regression Model accuracy metrics were carried out with the Root Mean Squared Error (RMSE), Mean Absolute Error (MAE) and R-squared. In terms of the interpretation of RMSE and MAE values, phase error is deemed functional at values below 1.0 and below 10 % for foliation. Acceptable error for operational usage is yet to be determined, however error below 0.5 for phase and 5 % foliation is similar, if not better than errors resulting from human observations. Here it should be noted that the phenological validation data could potentially have error especially when various technicians are making observations and flight missions do not occur on the same day as ground observations which was the case for the 2019 dataset. Descriptive statistics and linear trend lines were used to perform an initial overview of the phenological and meteorological data available at the Britz research station.

## 3 Results and Discussion

### 3.1 Phenological Data Historical Overview

Phenological observations have been carried out at the Britz research station by the same technician between 2006 and 2019. Spring leafing out observations began typically before budburst when buds began swelling (phase 0.5). Regular visits to the plots were then realised, often even on a daily basis for the purpose of catching individual tree bud bursting events. Once leafing out occurs for all sampling trees, the observer visitation was limited to twice a week, and approximately once a week after phase 4.0. Fruiting and flowering events were typically recorded only when present and absent of magnitude. Figure 3 shows an overview of the yearly averaged spring phenological observations from 2006 to 2020 of the beech plot at the Britz research station.

**Figure 3:**
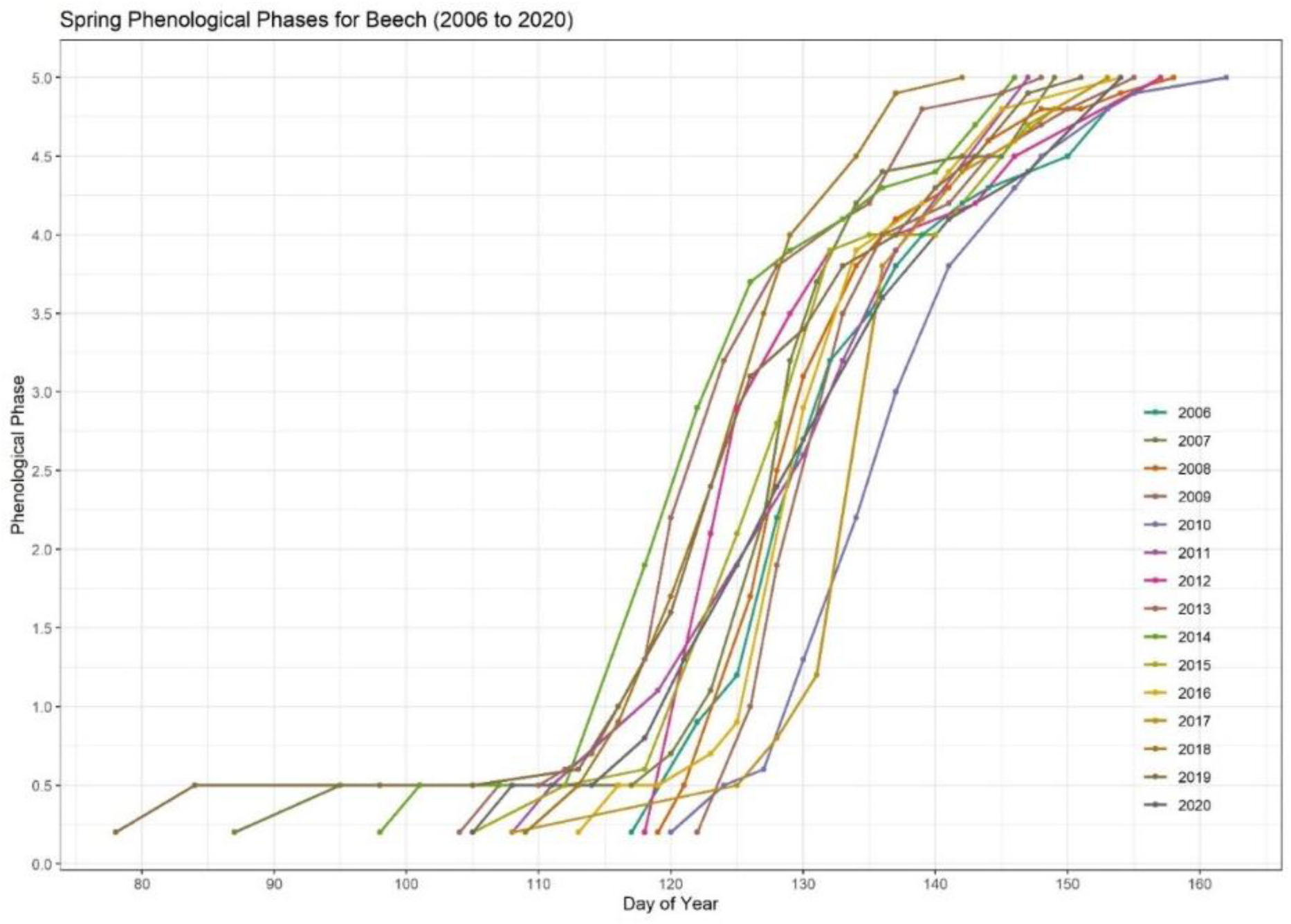
The phenological spring phase development for Beech at the Britz research station between 2006 and 2020.

The analysis of the time between phases is relevant for one, timing of budburst in terms of the effects of climate change, and two the arrival of the later phases such as phase 4 and phase 5 when leaves are close to being fully developed. The “hardening” of the leaf cell tissues is the morphological state where leaves will be less susceptible to late frost, possible early spring intense solar irradiation, as well as biotic pests (i.e. *Orchestes fagi*). During drought episodes in early spring, it could also be possible that specific phases are delayed and that the full development stages from phases 1.0 to 5.0 are prolonged. This may have been the case at the Britz research station in the years 2006, 2012, 2015, and 2019. Figure 4 gives a graphic representation of the length of the phases from 2006 to 2020 which varied from 23 to 41 days while Table 5 presents an overview with descriptive statistics for the length in days between phases. The possibility to predict other phases from a single phase observation would be a useful task especially for determining the dates for important “missed” phases at external plots. The phase length depicted in Figure 4 and Table 5 are based on the average phase timings for all of the beech plot phenology sampling trees. For the purpose of predicting other phases based on a single phase could possibly be modelled more appropriately using single tree data information as the heterogeneity among individuals can be very high during the spring phenological phases. Further research along these lines is warranted.

**Figure 4:**
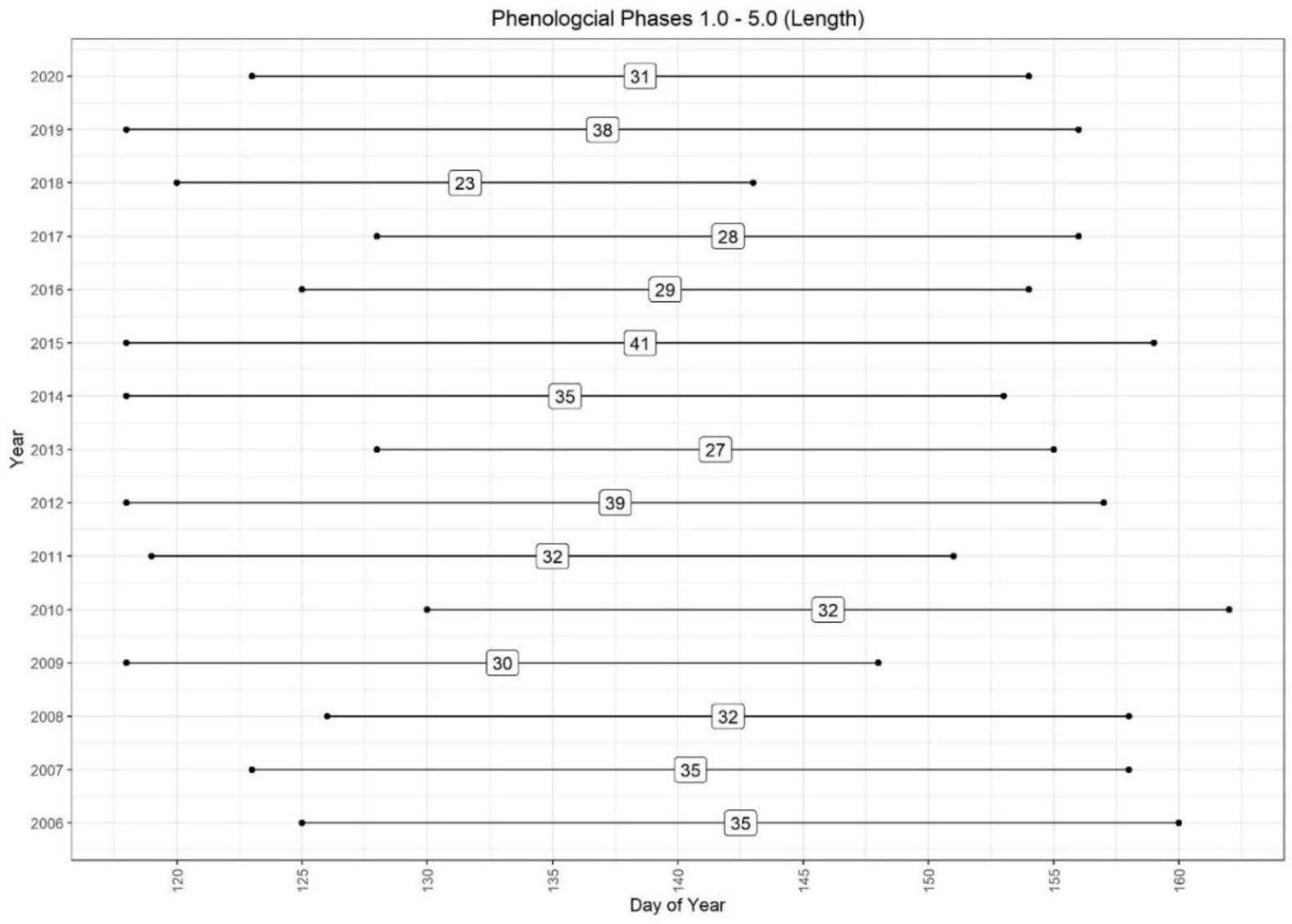
The average spring phenological phases at the Britz research station shown in length between phase 1 and 5 from years 2006 to 2020.

**Table 5:** An overview of the length between phases from 2006 to 2020. Accuracy is dependent on the temporal resolution of observations.

|  | 2006 | 2007 | 2008 | 2009 | 2010 | 2011 | 2012 | 2013 | 2014 | 2015 | 2016 | 2017 | 2018 | 2019 | 2020 | Min | Max | Mean | SD |
| --- | --- | --- | --- | --- | --- | --- | --- | --- | --- | --- | --- | --- | --- | --- | --- | --- | --- | --- | --- |
| Phase 1-2 | 4 | 4 | 2 | 2 | 4 | 11 | 6 | 5 | 4 | 8 | 5 | 3 | 3 | 5 | 6 | 2 | 11 | 4.8 | 2.3 |
| Phase 2-3 | 6 | 3 | 4 | 6 | 5 | 5 | 8 | 3 | 5 | 7 | 4 | 5 | 5 | 8 | 7 | 3 | 8 | 5.4 | 1.6 |
| Phase 3-4 | 14 | 13 | 12 | 12 | 11 | 12 | 10 | 9 | 12 | 13 | 8 | 8 | 7 | 14 | 8 | 7 | 14 | 10.9 | 2.4 |
| Phase 4-5 | 11 | 15 | 14 | 10 | 12 | 4 | 15 | 10 | 14 | 13 | 11 | 12 | 8 | 11 | 10 | 4 | 15 | 11.3 | 2.9 |
| Phase 1-4 | 24 | 20 | 18 | 20 | 20 | 28 | 24 | 17 | 21 | 28 | 17 | 16 | 15 | 27 | 21 | 15 | 28 | 21.1 | 4.3 |
| Phase 1-5 | 35 | 35 | 32 | 30 | 32 | 32 | 39 | 27 | 35 | 41 | 29 | 28 | 23 | 38 | 31 | 23 | 41 | 32.4 | 4.9 |

With regard to trends at the research station since 2006, it can been seen that phase 1.0 (see Figure 5; left) and phase 5.0 (see Figure 5; right) are trending towards an earlier onset. A gradual increase in average yearly air temperature (see Figure 6; left) is also evident, alongside a steady decrease in yearly precipitation (Figure 6; right).

**Figure 5:**
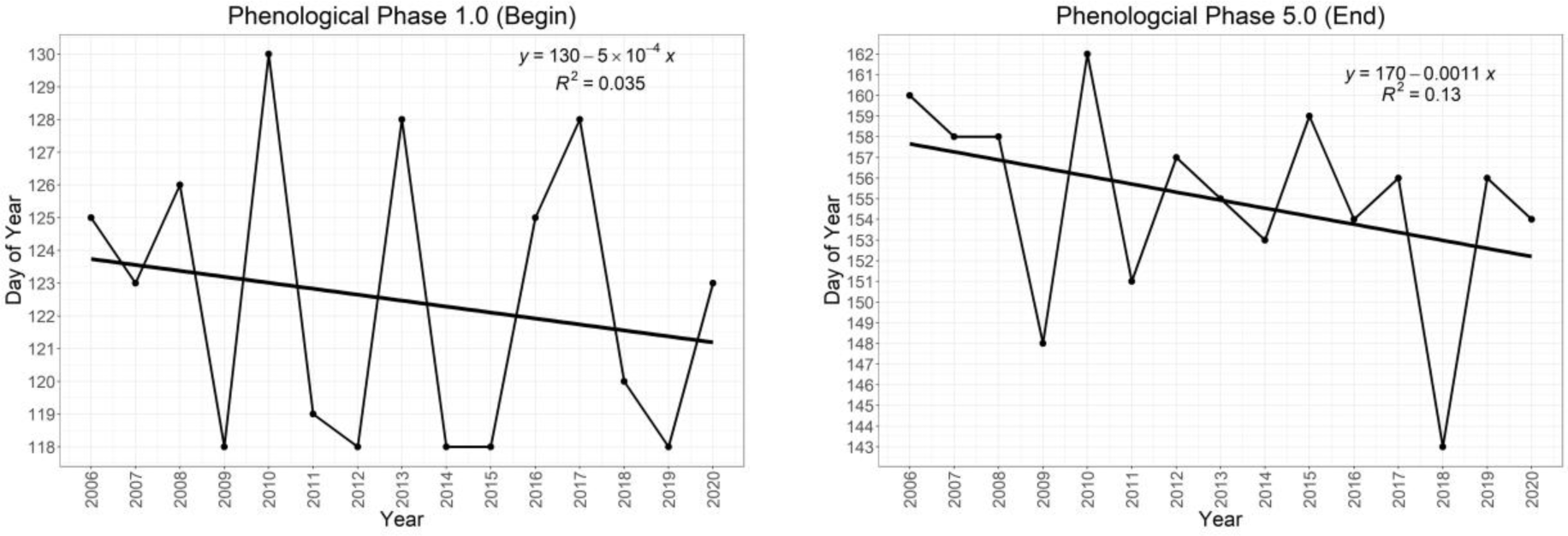
(left) Yearly linear trend the phenological phase 1.0; (right) Yearly linear trend of the phenological phase 5.0.

**Figure 6:**
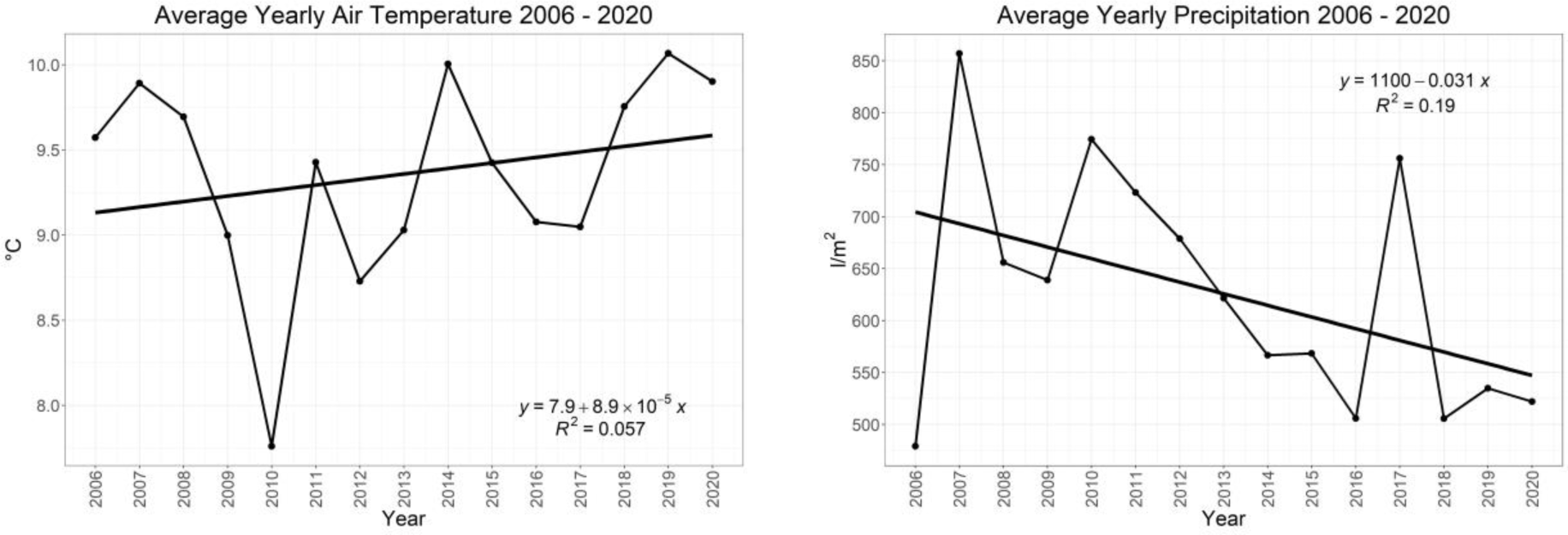
(left) Yearly linear trend of average air temperature between the years 2006 and 2020; (right) Yearly linear trend of average precipitation between the years 2006 and 2020. Both are results from the Britz research station.

Some of the phenological observation trees are equipped with electronic band dendrometers and sapflow measurements. Figure 7 shows the timing of the phenological phases for tree 328 in relation to the onset of stem growth at the beginning of the growth season. It can be seen in that in 2017 and 2018 the onset of stem diameter growth began around the same time as phase 3.0 was reached. Phase 3.0 coincides with the arrival of the first fully unfolded leaves. A significant periodical growth deficit fluctuations is evident in the 2018 dendrometer data throughout the growth season which coincides with the known long-lasting drought of that year (Schuldt et al., 2020).

**Figure 7:**
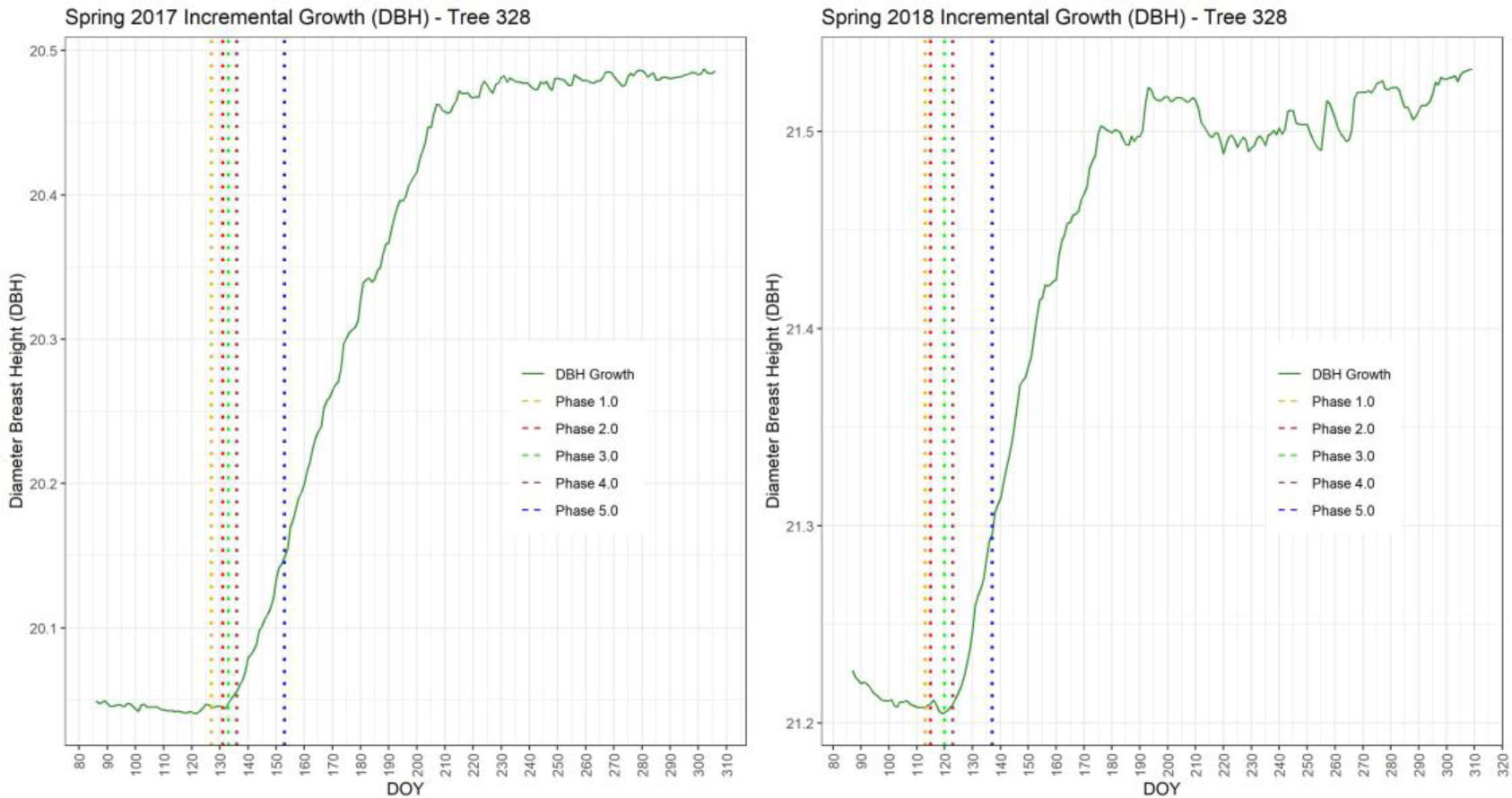
Spring phenological phases shown in relation to band dendrometer measurements from 2017 (left) and 2018 (right). Stem growth typically began around the arrival of phase 3.0.

With regard to an analysis into the phase and foliation datasets, the histograms shown in Figure 8 depict a clearly noticeable bimodal distribution, where the left and right skewed distributions are evident on the tail ends. This is caused by a typical surplus of observations before phase 1.0 due to increased observations in the anticipation of budburst and the length of time between phases 4.0 and 5.0.

**Figure 8:**
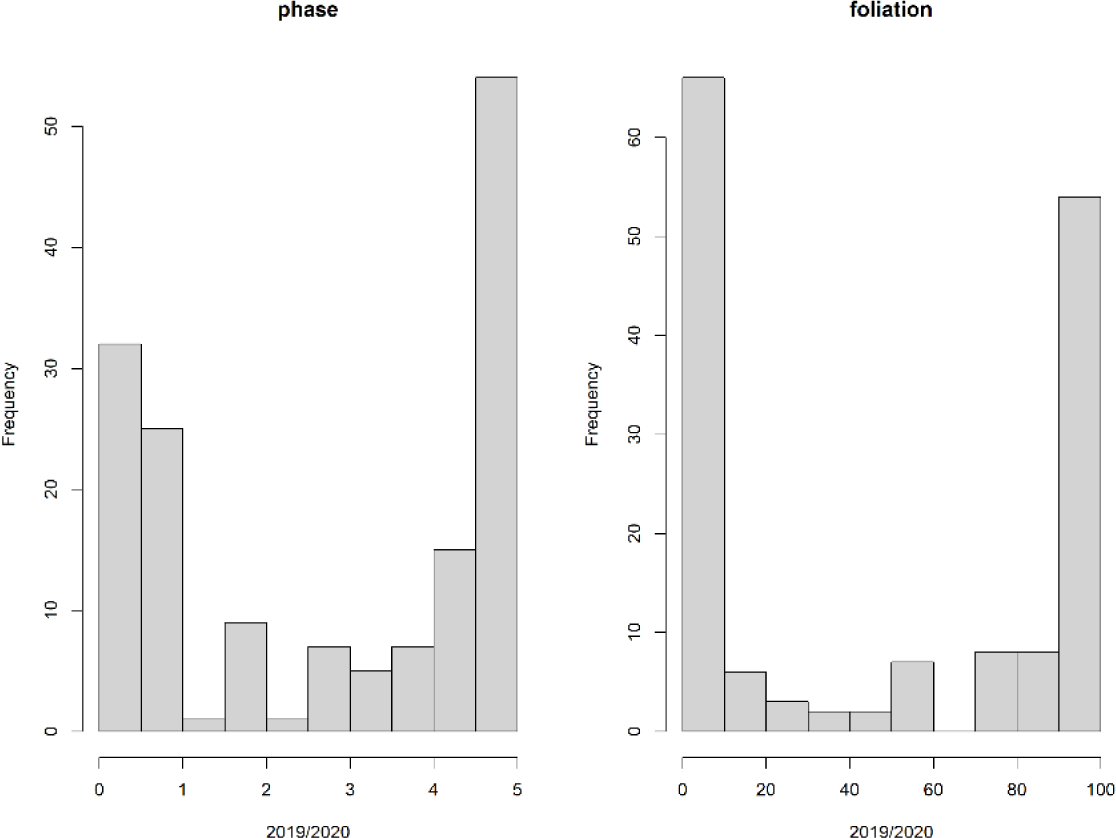
Histograms showing a distinct biomodial distribution of the phase and foliation ground observations from 2019 and 2020.

### 3.2 Correlation Analysis and Feature Selection

Due to the spectral reflectance characteristics of vegetation, visible bands tent to show a positive correlation among each other, whereas the NIR band a negative one (Mather & Koch, 2011). All of the vegetation indices whether derived from visible or NIR bands or a combination thereof, have a positive correlation with the phase and foliation datasets except for the NDWI which has typically an inverse relationship with the phases and foliation (see Figure 9). The most consistent index throughout all datasets, whether originating from single or combined years, is evidently the NDVI with a persistent correlation of *r* > 0.9 (p < 0.001) over all datasets.

**Figure 9:**
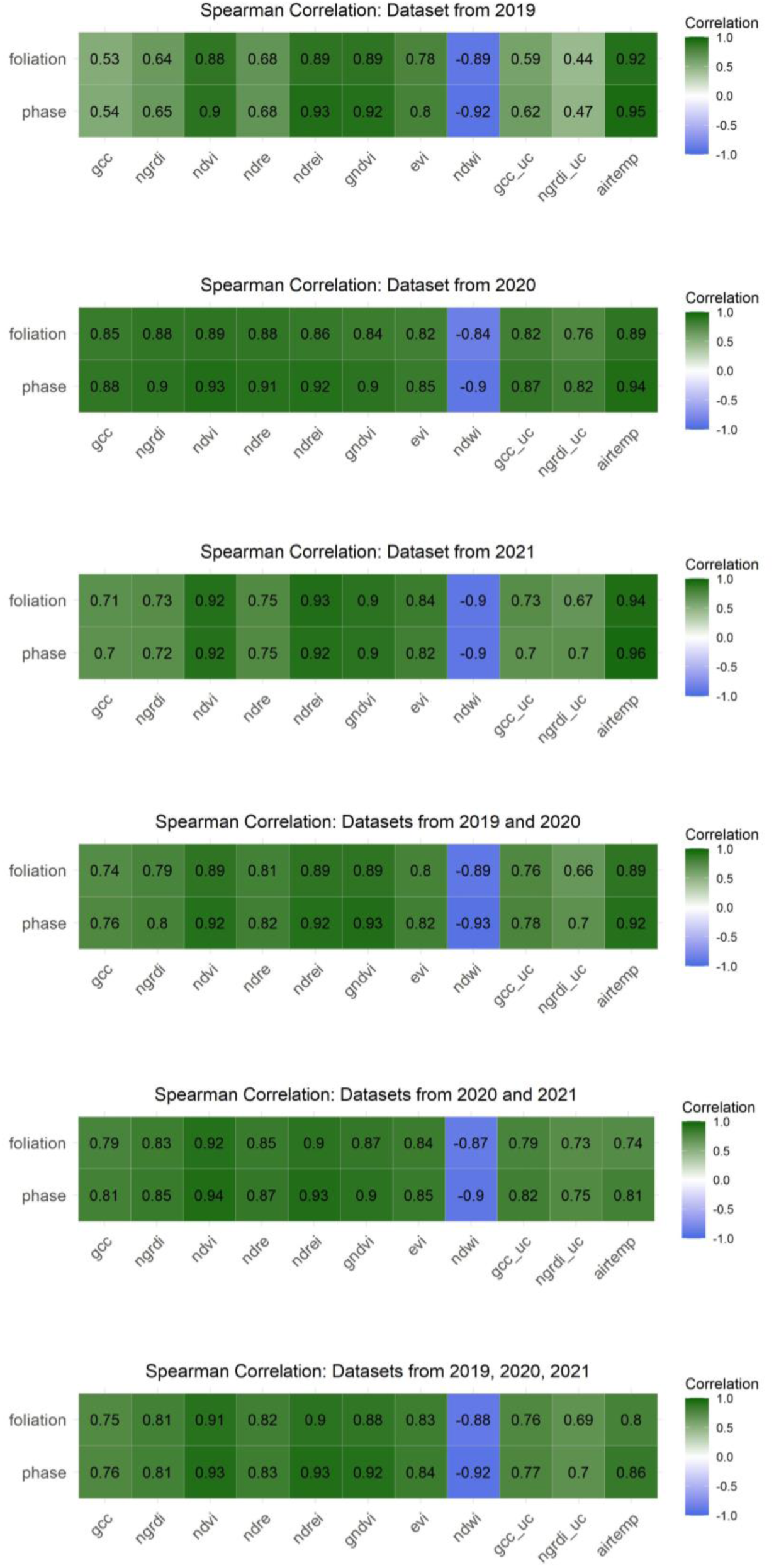
Spearman correlation analysis of the spectral indices derived from the 2019, 2020 and 2021 datasets in relation to the ground observations.

Indices derived from visual bands (i.e. GCC and NGRDI) showed a correlation of *r* = 0.65 (p < 0.001) and those uncalibrated even poorer. Interestingly, the AIRTEMP meteorological-based feature correlated very well with the ground observations (*r* = 0.9; p < 0.001) with a very high correlation coefficient to the phenological phases at *r* = 0.95 (p < 0.001).

In terms of correlation among independent features (see Figure 10), the aim was to refrain from implementing highly correlated features when multiple independent features where to be incorporated into the modelling process. This could be especially problematic when multiple indices are derived from the same bands (i.e. NDVI and EVI). Here we could deduce that the NDREI and GCC, when used together for the modelling process, have a lower correlation (*r =* 0.73) and do not share any similar bands. Likewise, the NDRE and the NDWI do not share the same bands and have a negative correlation coefficient of *r* = −0.8. The NDWI and the GCC share only the green band and correlate negatively at *r* = −0.74.

**Figure 10:**
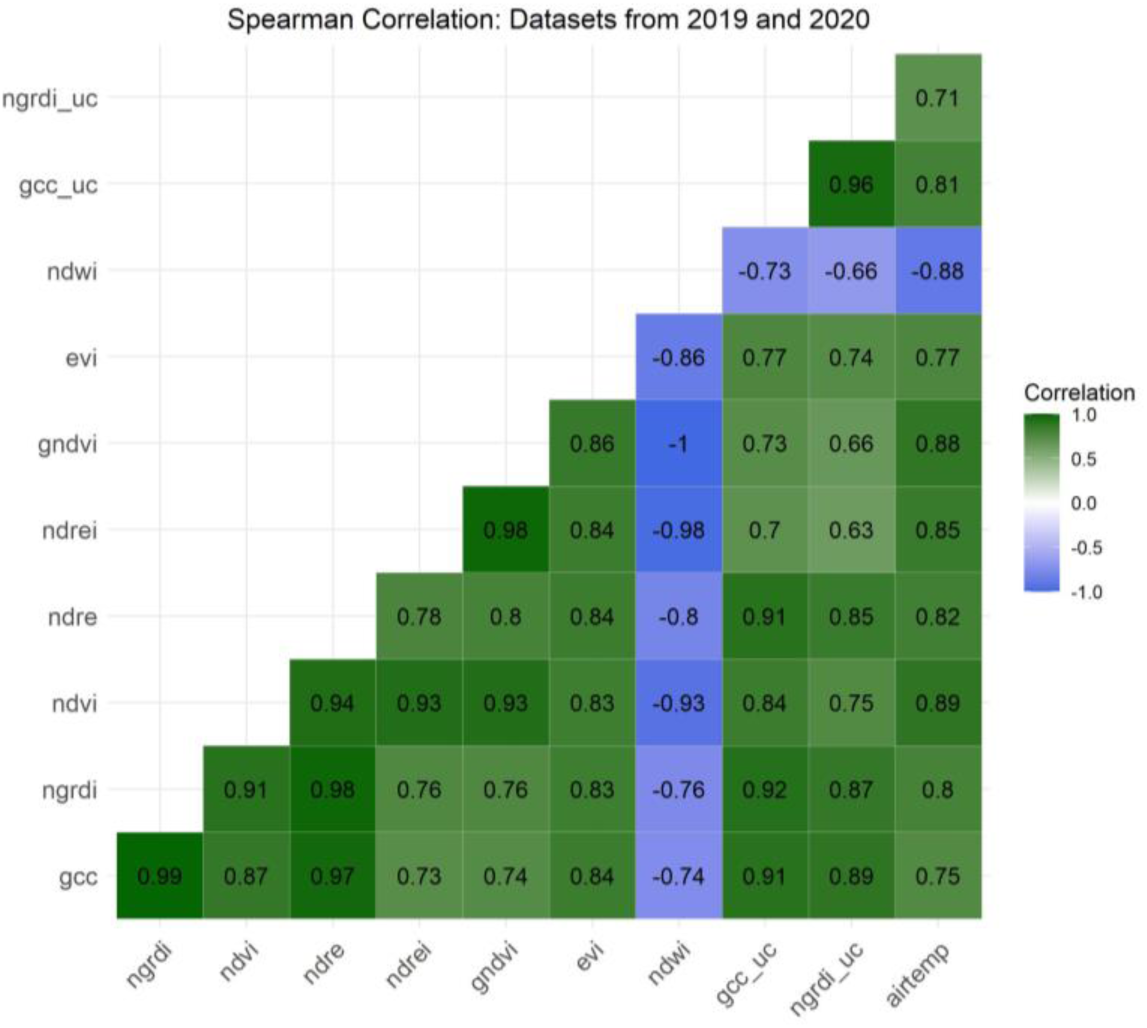
Between-variable Spearman correlation assessment of the 2019/2020 features.

The use of correlation for the purpose of feature selection, though informative especially for evaluating multicollinearity, could potentially be misleading as correlation coefficients will be high due to the bimodal influence on the dataset. Here the aggregation of data points on the tail ends will result in a biased similarity due to an oversampling of similar phases thus causing a high correlation coefficient. For this reason we did not rely solely on correlation filtering methods (Chandrashekar & Sahin, 2014) for a finalised feature selection.

### 3.3 Polynomial Regression and Feature Selection

The addition of polynomial terms into regression models can aid in characterising nonlinear patterns (Kuhn & Johnson, 2019) and is conducive to representing phenological trends in particular that of the spring green-up phases. As the polynomial fitting may not be capable in identifying the complexities of phenology metrics in comparison to other algorithms (Rodrigues et al., 2012; Zhu et al., 2012), we used the fitting of polynomials here for the purpose of feature selection, where the aim was to identify which features best correspond to the typical spring phenology curve. Figure 11 shows the fitting of the five polynomial orders using the example for the NDVI resulting in an RMSE 0.55, MAE of 0.41 and R-squared of 0.91. Here the third polynomial order was deemed the best choice for further analysis where the curve is neither oversimplified nor too complex.

**Figure 11:**
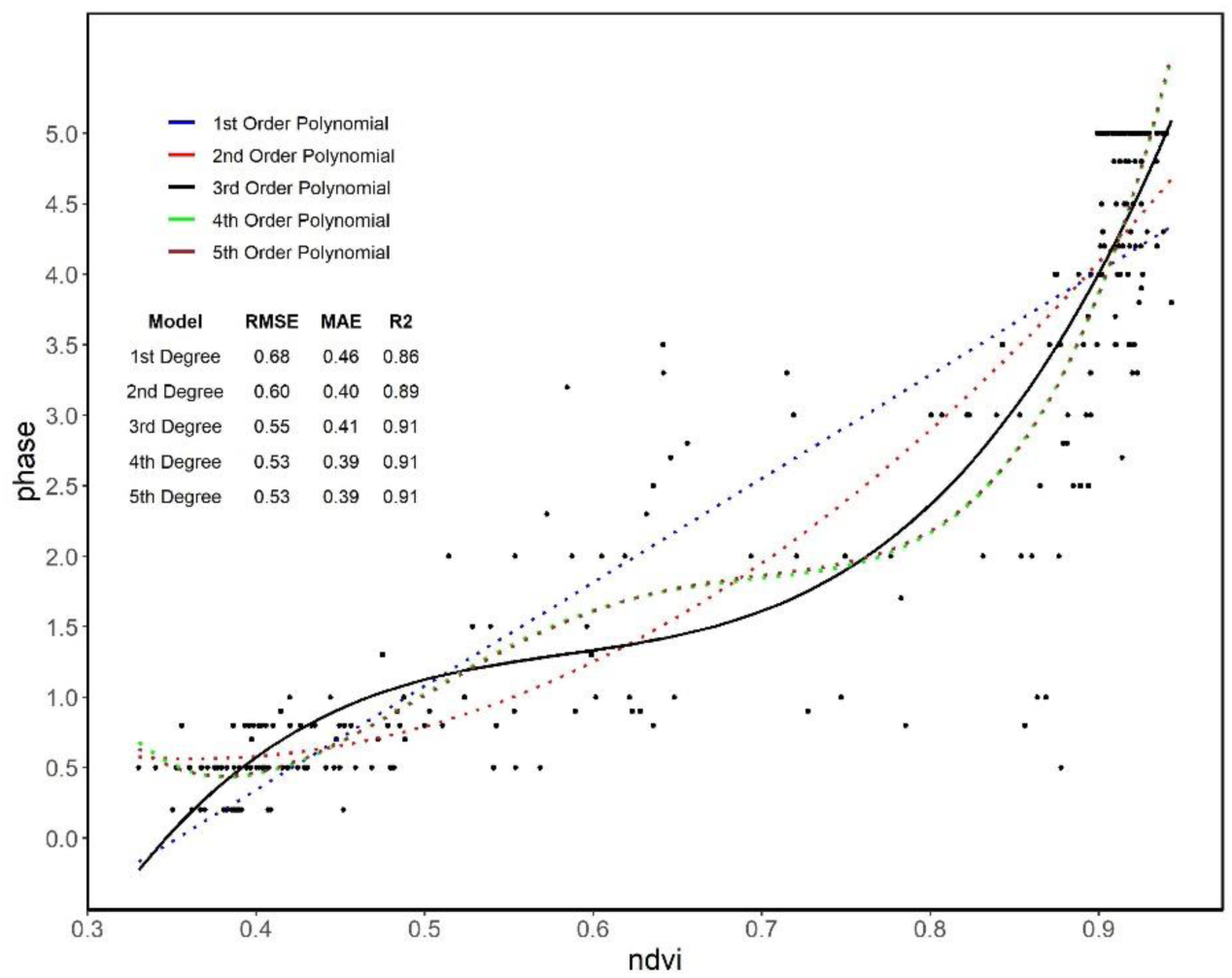
Modelling of the spring phenological phases (2019/2020) dataset with polynomial regression of the first to fifth order.

To follow, each of the selected individual features was tested with the 3^rd^ order polynomial separately for the 2019/2020 and 2020/2021 datasets for both phase (Figure 12) and foliation (Figure 13). In terms of the phenological phases, the GNDVI shows quite a low dispersal of RMSE for the 2019/2020 dataset yet the dispersal is higher for the 2020/2021 dataset. A similar result is evident for the NDVI where however less dispersal is found in the 2020/2021 dataset rather than that of the 2019/2020 dataset. The cumulative warming days (AIRTEMP) as well as

**Figure 12:**
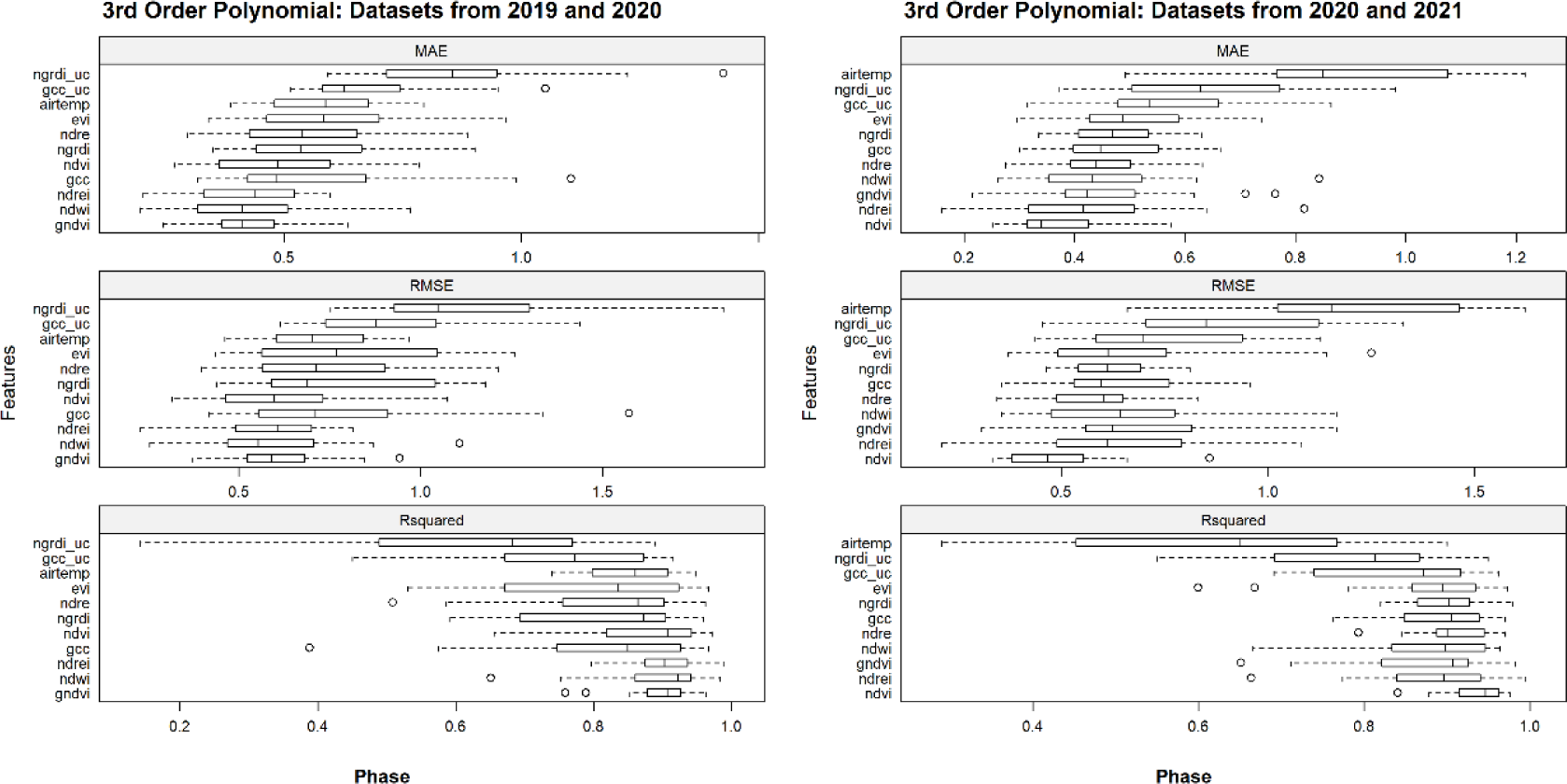
Overview of the spring phenological phases and indices modelled with polynomial regression of the third order for the 2019/2020 (left) and 2020/2021 (right) datasets.

**Figure 13:**
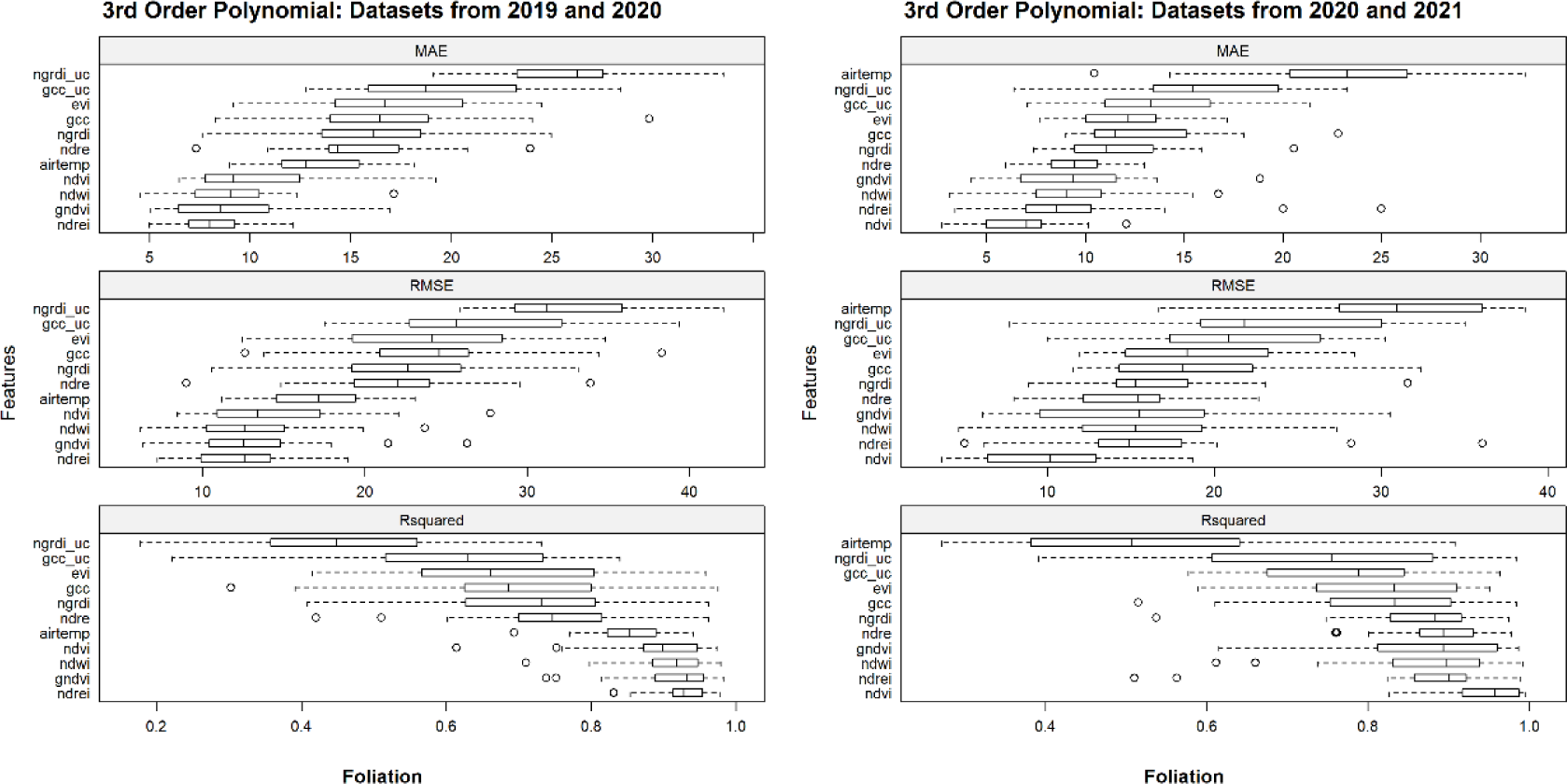
Overview of spring foliation and indices modelled with polynomial regression of the third order for the 2019/2020 (left) and 2020/2021 (right) datasets.

the indices derived from the uncalibrated visible bands (GCC_UC and NGRDI_UC) fared poorly for both datasets. This was also the case for foliation, however AIRTEMP performed better for the 2019/2020 dataset. With regard to foliation, the NDVI also performed well for the 2020/2021 dataset as did the NDREI for both datasets.

### 3.4 Machine Learning Models: 2019/2020 Datasets

Based on the results of the correlation analysis and polynomial fitting, we were able to select the most relevant features for further scrutinisation during the subsequent modelling process. Important to note here is that in the initial feature selection process using only the correlation analysis alone could have produced an unseen bias due to an aggregation of data points at the tail ends of the datasets which was evident especially for the 2019/2020 dataset. We proceeded to build three models based on ML algorithms which in effect aided in choosing the best performing algorithms as well as features. Each of the selected individual and combined indices were modelled with each algorithm and evaluated using an 80/20 training/validation data split. This not only helped in choosing the best ML algorithm, but also assisted in a type of model-based feature selection by further narrowing down the selected features. In terms of the phenological phases, an RMSE of ≤ 0.5 (0.6) is deemed as acceptable and similar to the magnitude of potential human error. For the Britz method of foliation, a RMSE of ≤ 10 % is assumed to be acceptable, however some may argue that a RMSE of ≤ 5 % in terms of foliage observations is possible with ground observation. Here it should be reminded that the Britz method of foliation is based on the percentage of leaves which have fully opened rather than fractional cover or greening-up.

#### Phenological Phases

With regard to the phenological phases, the GAM Boosting algorithm showed overall the best results (see Table 6). The GAM models with the features NDREI + GCC resulted with a RMSE of 0.51, MAE of 0.33 and an R-squared of 0.95. The feature combination of NDWI + GCC resulted with a RMSE of 0.46, MAE of 0.3 and R-squared of 0.96. The top performing model was that of the GAM Boosting with the NDVI and produced a RMSE of 0.28, MAE of 0.18, and R-squared of 0.98. The second-best performing model was that of the GAM model with the NDRE + NDWI input features resulting with a RMSE of 0.44, MAE of 0.31 and R-squared of 0.96. Interestingly, the uncalibrated GCC (GCC_UC) outperformed the calibrated GCC with a RMSE of 0.73 for Gradient Boosting and the GCC_UC index as opposed to a RMSE of 0.81 for GAM Boosting and the GCC.

**Table 6:**
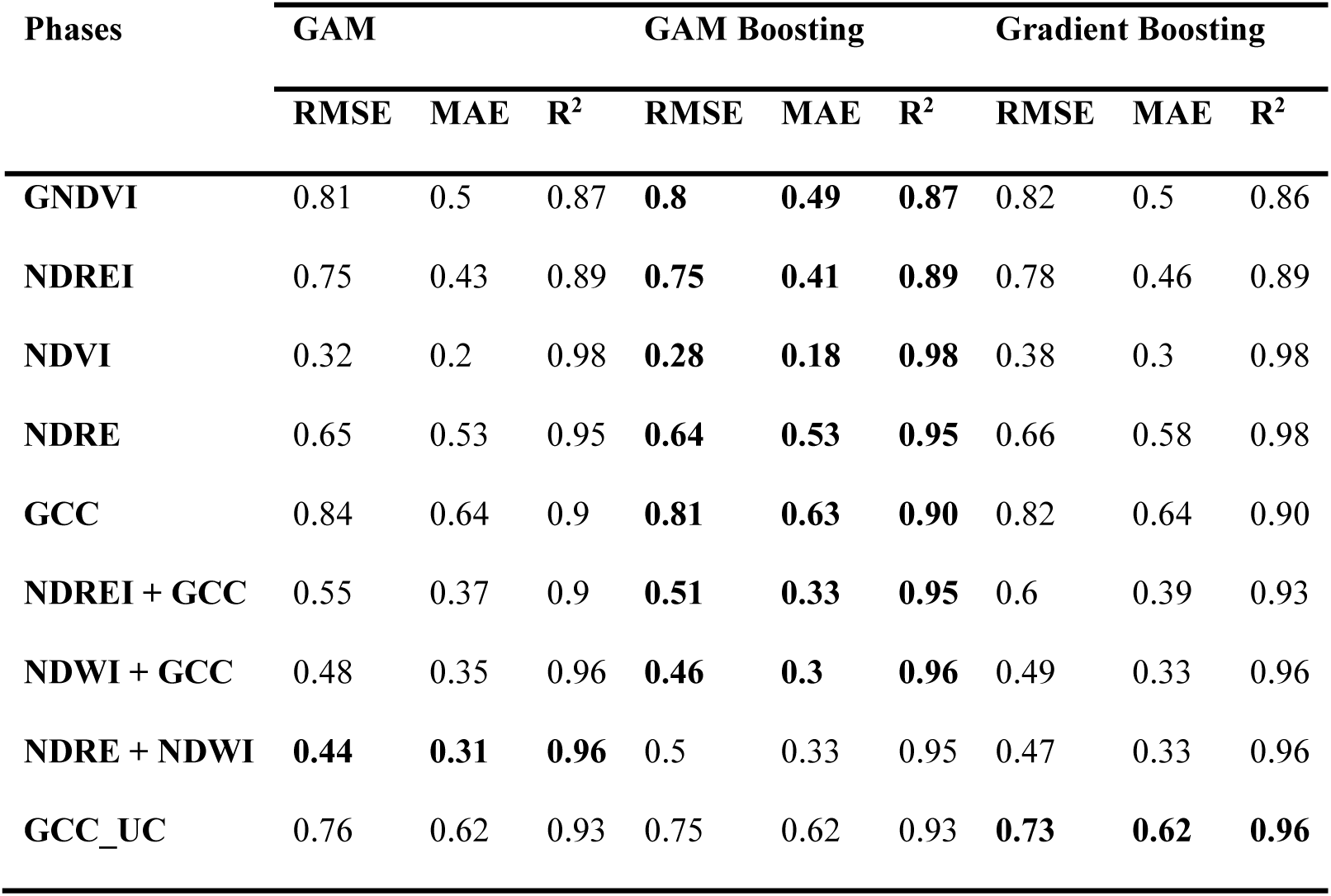
Error metrics for the phase prediction of three Machine Learning algorithms. Values shown in bold font depict the best results.

| Phases | GAM |  |  | GAM Boosting |  |  | Gradient Boosting |  |  |
| --- | --- | --- | --- | --- | --- | --- | --- | --- | --- |
|  | RMSE | MAE | R<sup>2</sup> | RMSE | MAE | R<sup>2</sup> | RMSE | MAE | R<sup>2</sup> |
| **GNDVI** | 0.81 | 0.5 | 0.87 | **0.8** | **0.49** | **0.87** | 0.82 | 0.5 | 0.86 |
| **NDREI** | 0.75 | 0.43 | 0.89 | **0.75** | **0.41** | **0.89** | 0.78 | 0.46 | 0.89 |
| **NDVI** | 0.32 | 0.2 | 0.98 | **0.28** | **0.18** | **0.98** | 0.38 | 0.3 | 0.98 |
| **NDRE** | 0.65 | 0.53 | 0.95 | **0.64** | **0.53** | **0.95** | 0.66 | 0.58 | 0.98 |
| **GCC** | 0.84 | 0.64 | 0.9 | **0.81** | **0.63** | **0.90** | 0.82 | 0.64 | 0.90 |
| **NDREI + GCC** | 0.55 | 0.37 | 0.9 | **0.51** | **0.33** | **0.95** | 0.6 | 0.39 | 0.93 |
| **NDWI + GCC** | 0.48 | 0.35 | 0.96 | **0.46** | **0.3** | **0.96** | 0.49 | 0.33 | 0.96 |
| **NDRE + NDWI** | **0.44** | **0.31** | **0.96** | 0.5 | 0.33 | 0.95 | 0.47 | 0.33 | 0.96 |
| **GCC_UC** | 0.76 | 0.62 | 0.93 | 0.75 | 0.62 | 0.93 | **0.73** | **0.62** | **0.96** |

At this stage of the modelling process, the NDVI and GAM boosting algorithm showed very good results (RMSE = 0.28) and the question is here whether the dataset is overfit for the Britz research station beech stand. At this point it is an imperative to test the models with unseen data and assess which ones are generalizable over various Beech stands especially those of increased age. In terms of the models derived from indices from the visual bands, the uncalibrated GCC performed slightly better than that of the radiometrically calibrated one and better than some of the models derived from the calibrated multispectral bands which is particularly interesting as RGB sensors are typically acquired at a much cheaper price.

#### Foliation

For the most part, all models failed the 10 % cutoff point except for those using the NDVI as an input feature. Both the NDVI-based GAM boosting and Gradient Boosting models obtained a RMSE of 7 %, MAE of 4 % and R-Squared of 0.98. Here overfitting could also be factor, however will still be interesting for further model assessment of the prediction of foliation on a new dataset (2022) as well as datasets outside of the Britz research station. The worst performing models were those utilising the radiometrically calibrated GCC which acquired a RMSE of 22 %, MAE of 16 %, and R-squared of 0.92.

### 3.5 GAM Boosting Models with Test Datasets

With the aim of testing the robustness as well as generalisability of the developed models, new data from 2022 as well as data from different forest stands (beech) was introduced. Here we tested the models on new spring phenological data from the same stand from 2022 (*n* = 17) as well as older beech stand in Kahlenberg (*n* = 10) located in the same region as the Britz research station and a beech stand in the more mountainous region of the Black Forest (*n* = 8) in south-western Germany. The three test datasets are limited to only one Epoch where the Kahlenberg site is comprised of mostly later phases and the Britz and Black Forest datasets have a wide range of earlier phases (< 4.0). Additionally, training datasets were divided into three different subdivisions based on the year of origin: 2019/2020, 2020/2021 and all datasets together (2019 – 2021). This was carried out for the purpose of distinguishing whether data acquisition methods from a certain year contributed to error propagation. For example, the 2019 field data was carried out by a different observer and often not recorded on the same day as flights (± 3 days) as well as low quality radiometric calibration. The models chosen for testing were those implementing GAM boosting and the RGB-derived indices GCC (Micasense Altum) and GCC_UC (Zenmuse X7) and the NDVI (Micasense Altum). Table 8 displays a list of all of the tested models with reference to the applied index, location, training data subdivision and date.

**Table 7:**
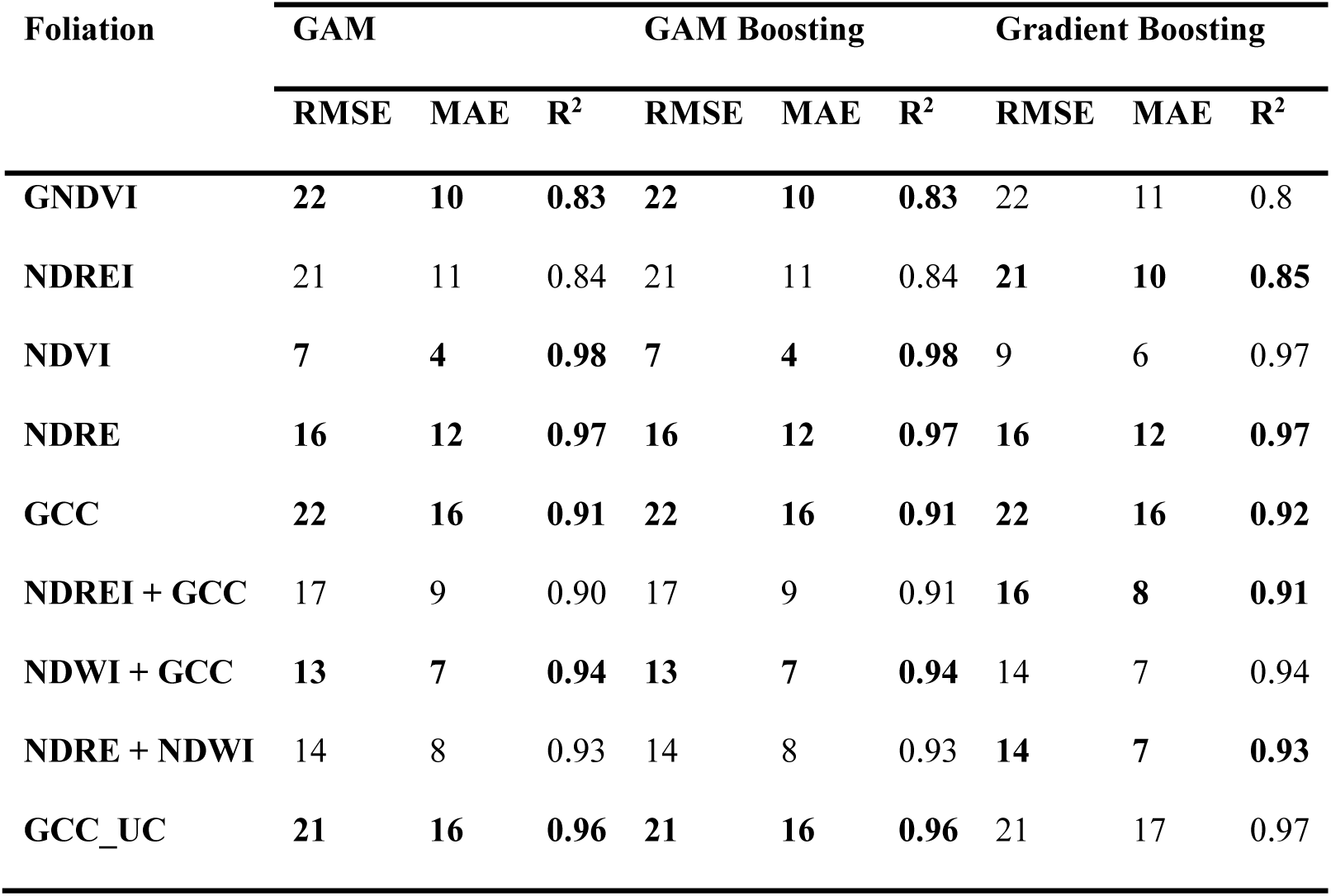
Error metrics (in %) for the foliation prediction of three Machine Learning algorithms. Values shown in bold font depict the best results.

| Foliation | GAM |  |  | GAM Boosting |  |  | Gradient Boosting |  |  |
| --- | --- | --- | --- | --- | --- | --- | --- | --- | --- |
|  | RMSE | MAE | R<sup>2</sup> | RMSE | MAE | R<sup>2</sup> | RMSE | MAE | R<sup>2</sup> |
| **GNDVI** | **22** | **10** | **0.83** | **22** | **10** | **0.83** | 22 | 11 | 0.8 |
| **NDREI** | 21 | 11 | 0.84 | 21 | 11 | 0.84 | **21** | **10** | **0.85** |
| **NDVI** | **7** | **4** | **0.98** | **7** | **4** | **0.98** | 9 | 6 | 0.97 |
| **NDRE** | **16** | **12** | **0.97** | **16** | **12** | **0.97** | **16** | **12** | **0.97** |
| **GCC** | **22** | **16** | **0.91** | **22** | **16** | **0.91** | **22** | **16** | **0.92** |
| **NDREI + GCC** | 17 | 9 | 0.90 | 17 | 9 | 0.91 | **16** | **8** | **0.91** |
| **NDWI + GCC** | **13** | **7** | **0.94** | **13** | **7** | **0.94** | 14 | 7 | 0.94 |
| **NDRE + NDWI** | 14 | 8 | 0.93 | 14 | 8 | 0.93 | **14** | **7** | **0.93** |
| **GCC_UC** | **21** | **16** | **0.96** | **21** | **16** | **0.96** | 21 | 17 | 0.97 |

**Table 8:** displays the various models tested on the datasets from 2022 and/or outside of the Britz research station. The four models in bold font are those deemed operational.

| Model Name | Site | Index | Training | DOY | Year |
| --- | --- | --- | --- | --- | --- |
| br-ndvi-19-20 | Britz | ndvi | 2019-2020 | 125 | 2022 |
| **br-ndvi-20-21** | **Britz** | **ndvi** | **2020-2021** | **125** | **2022** |
| br-ndvi-19-21 | Britz | ndvi | 2019-2021 | 125 | 2022 |
| br-gcc-19-20 | Britz | gcc | 2019-2020 | 125 | 2022 |
| **br-gcc-20-21** | **Britz** | **gcc** | **2020-2021** | **125** | **2022** |
| br-gcc-19-21 | Britz | gcc | 2019-2021 | 125 | 2022 |
| br-gcc-uc-19-20 | Britz | gcc_uc | 2019-2020 | 125 | 2022 |
| br-gcc-uc-20-21 | Britz | gcc_uc | 2020-2021 | 125 | 2022 |
| br-gcc-uc-19-21 | Britz | gcc_uc | 2019-2021 | 125 | 2022 |
| **bf-gcc-19-20** | **Black Forest** | **gcc** | **2019-2020** | **115** | **2022** |
| bf-gcc-20-21 | Black Forest | gcc | 2020-2021 | 115 | 2022 |
| bf-gcc-19-21 | Black Forest | gcc | 2019-2021 | 115 | 2022 |
| bf-gcc-uc-19-20 | Black Forest | gcc_uc | 2019-2020 | 115 | 2022 |
| bf-gcc-uc-20-21 | Black Forest | gcc_uc | 2020-2021 | 115 | 2022 |
| bf-gcc-uc-19-21 | Black Forest | gcc_uc | 2019-2021 | 115 | 2022 |
| ka-ndvi-19-20 | Kahlenberg | ndvi | 2019-2020 | 119 | 2020 |
| ka-ndvi-20-21 | Kahlenberg | ndvi | 2020-2021 | 119 | 2020 |
| ka-ndvi-19-21 | Kahlenberg | ndvi | 2019-2021 | 119 | 2020 |
| ka-gcc-19-20 | Kahlenberg | gcc | 2019-2020 | 135 | 2020 |
| ka-gcc-20-21 | Kahlenberg | gcc | 2020-2021 | 135 | 2020 |
| ka-gcc-19-21 | Kahlenberg | gcc | 2019-2021 | 135 | 2020 |
| ka-gcc-uc-19-20 | Kahlenberg | gcc_uc | 2019-2020 | 135 | 2020 |
| **ka-gcc-uc-20-21** | **Kahlenberg** | **gcc_uc** | **2020-2021** | **135** | **2020** |
| ka-gcc-uc-19-21 | Kahlenberg | gcc_uc | 2019-2021 | 135 | 2020 |

Results of the model testing of the phenological phase prediction (see Figure 14) and foliation (see Figure 15) were ranked in order of the RMSE. Noteworthy is that all the models of the phenological phase prediction that achieved the 0.5 threshold (left of green dotted line) were those of the calibrated and uncalibrated GCC which originate from bands of the visible portion of the electromagnetic spectrum. Five of six of these models were from the Kahlenberg dataset and one from the Black Forest dataset. The best performing models where selected for each of the test sites and are mapped out in Figures 16-19. All image data acquired for the test sites with the *Zenmuse X7* lack radiometric calibration except for the Britz dataset (see Figure 19) which was acquired with the with both the *X7* and radiometrically calibrated *Micasense Altum* data.

**Figure 14:**
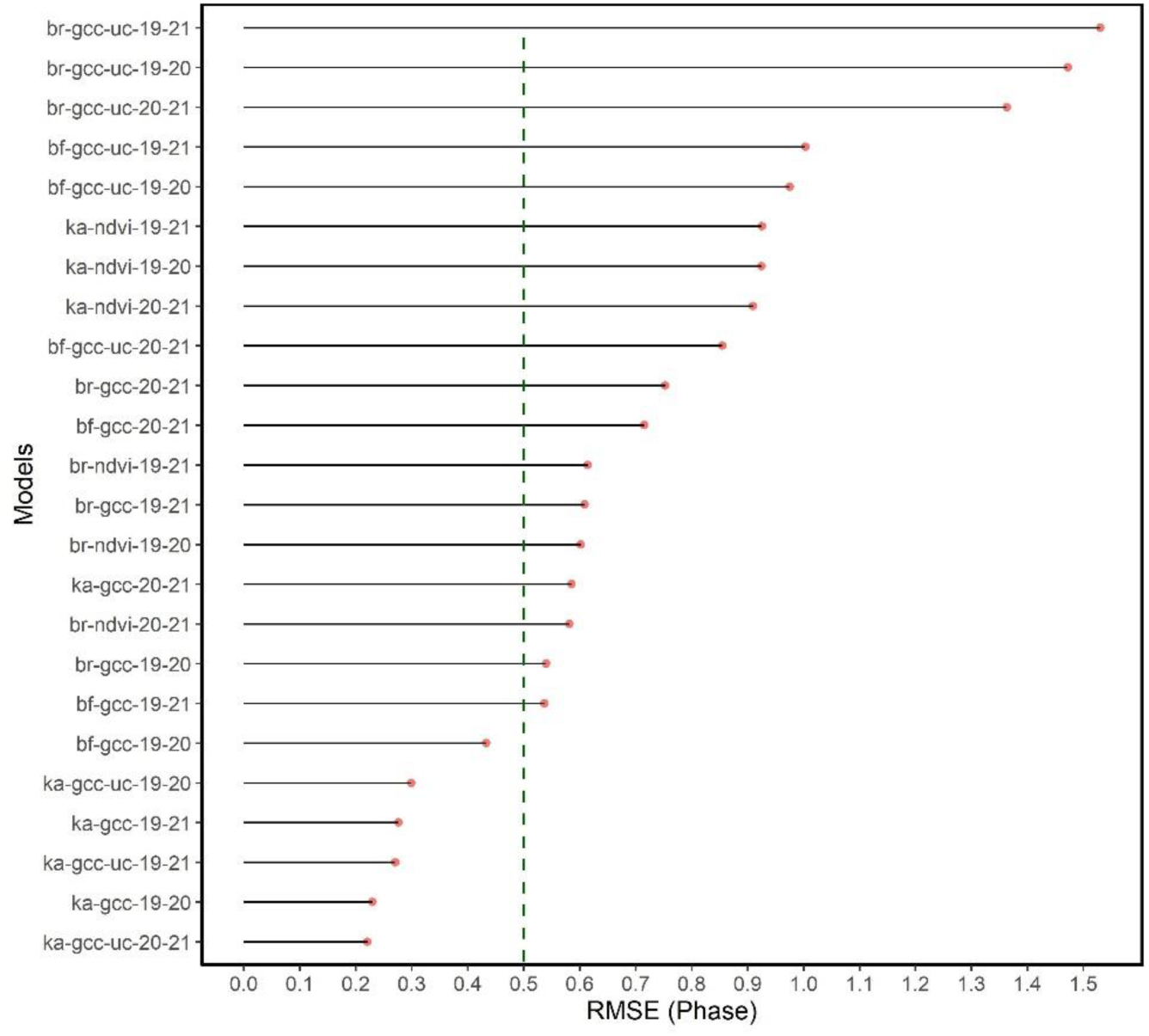
graph showing the RMSE for the phase prediction ranked in order from poorest to best RMSE. The green dashed line depicts the cut-off point of acceptable accuracy. Allowing an RMSE of up to 0.6 would enable the NDVI model derived from the multispectral datasets. Otherwise, only models originating from the visible bands are considered operational.

**Figure 15:**
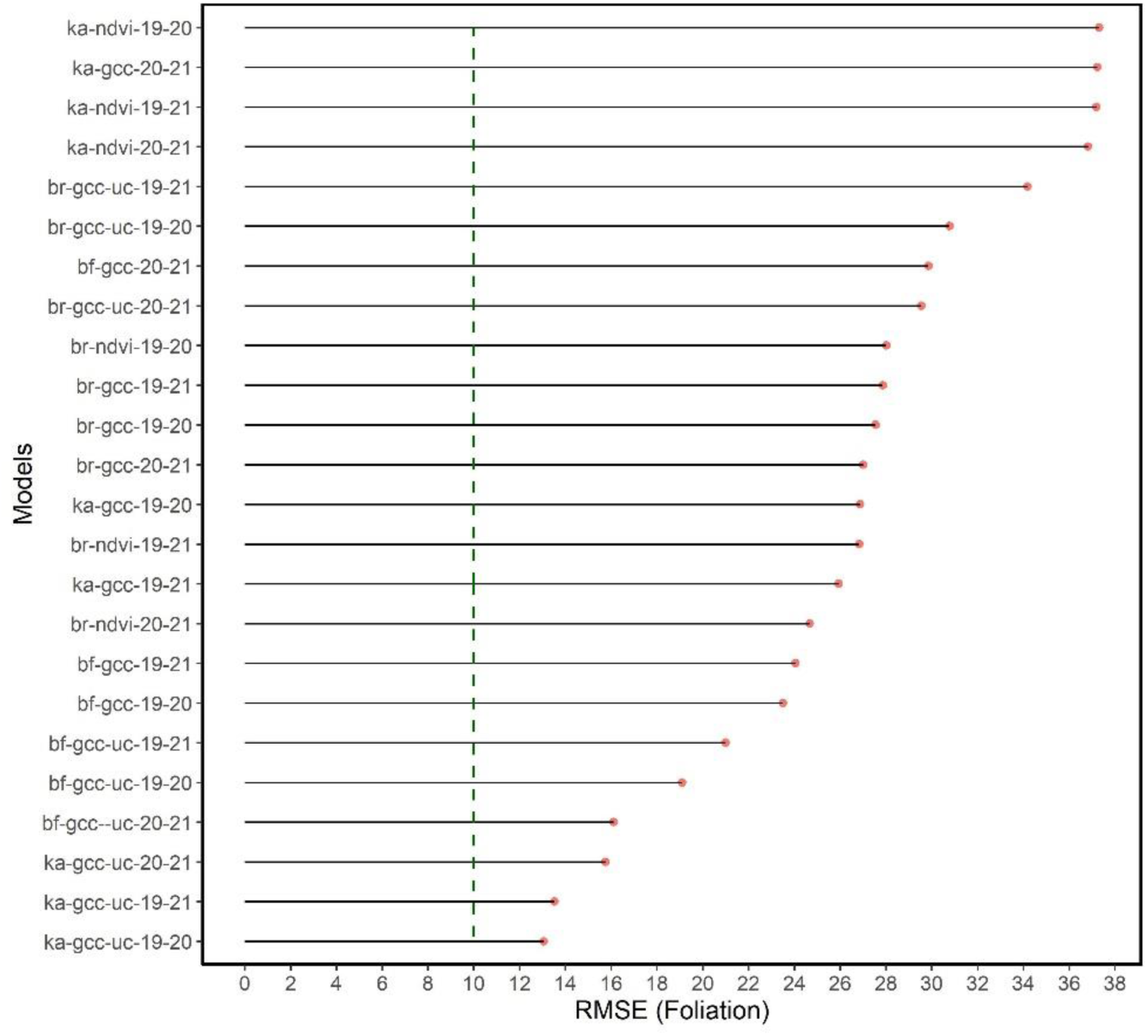
graph showing the RMSE for foliation prediction ranked in order from poorest to best. The green dashed line depicts the cut-off point of 10 %. None of the models for foliation prediction are considered functional.

**Figure 16:**
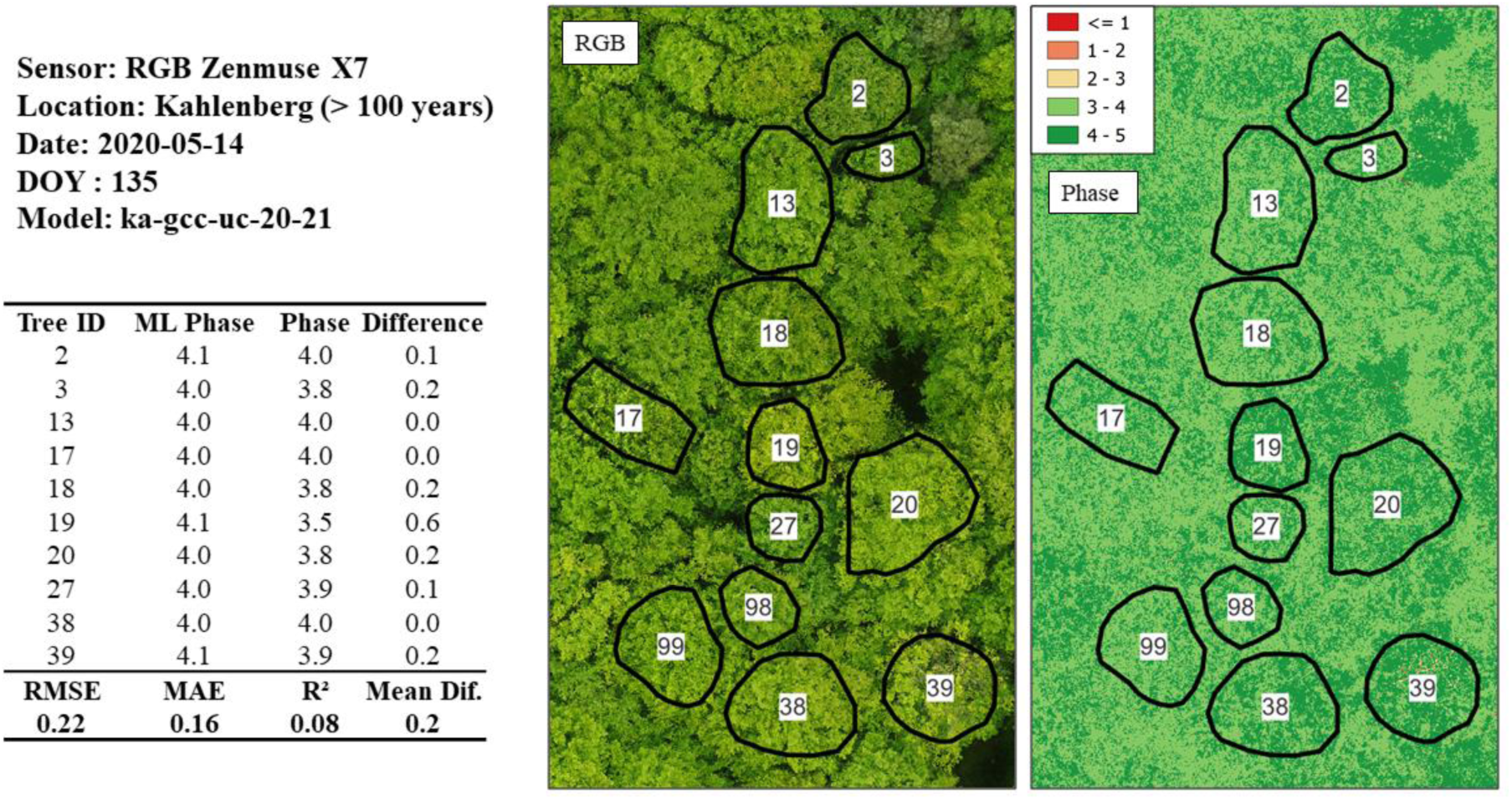
Phase prediction of an older Beech stand (> 100 years) utilising the model originating from the uncalibrated GCC 2020/2021 dataset. The very low RMSE of 0.22 proves a highly generalizable model however it should be noted that this is a relatively small dataset (n = 10) and comprised of only later phases (> 3.0). “ML phase” is the predicted phase and “Phase” originates from the ground-based observations.

The Kahlenberg dataset (see Figure 16) with the *gcc-uc-20-21* model resulted with a very low RMSE of 0.22, MAE of 0.16 and R-Squared of 0.08 (*n* = 10). Such a low RMSE for an uncalibrated RGB-based model is an unexpected result here and shows that the later phases, in particular phase 4.0 predict well. Phase 4.0 is a significant phase in the spring green-up as it corresponds to the completion of all leaf and shoot development. The transition to Phase 5.0 would then follow with the hardening of leaf tissue alongside a change to darker green and increased late-frost hardiness.

With regard to the Black Forest dataset with the *bf-gcc-19-20* model, a RMSE of 0.43, MAE of 0.32, and R-squared of 0.02 (*n* = 8) was achieved (see Figure 17). Here, a scene with a wide range of phases (0.9 – 3.8) was available, and a successful phenological phase prediction was possible with the calibrated GCC model and training data from 2019 and 2020. Important to note is that the radiometrically calibrated GCC model was used to predict on the GCC which is derived from the non-calibrated *Zenmuse X7*. Significant here is that sensor mixing in terms of model training with the multispectral sensor and prediction with a consumer grade RGB sensor is attainable. We considered the low R-Squared as insignificant due to the overall low sample rate of the test datasets.

**Figure 17:**
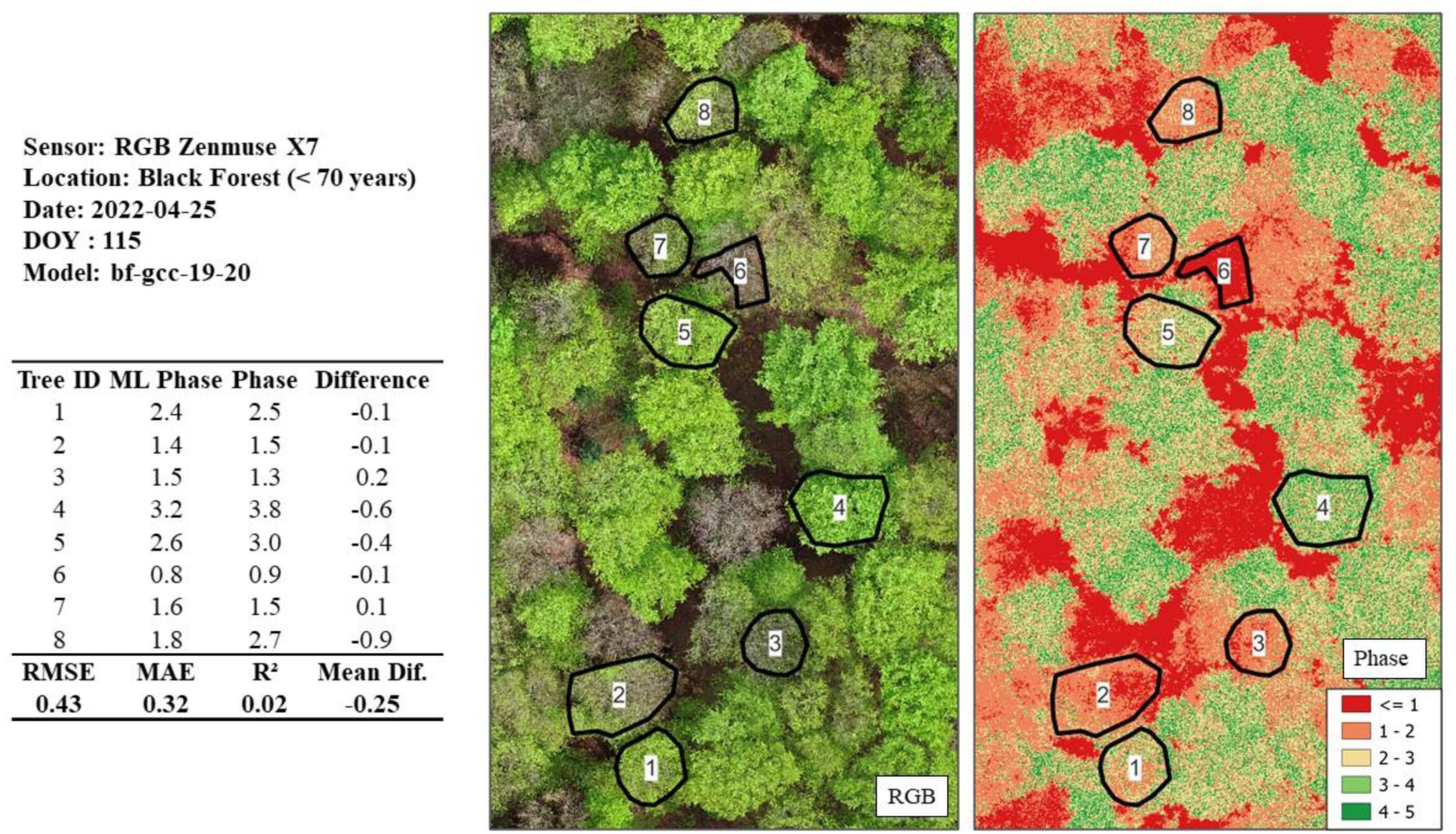
Phase prediction of a Beech stand (< 70 years) utilising the model originating from the calibrated GCC 2019/2020 dataset. The Black Forest dataset is a particularly challenging one as a wide range of phases are available. An RMSE of 0.43 is within the accepted error cut-off of ≤ 0.5.

The Britz dataset (see Figure 18) also implemented the GCC and 2019/2020 training model (*br- gcc-19-20)* and resulted in a RMSE of 0.54, MAE of 0.45 and R-squared of 0.65 (*n* = 17). Important to note is that the Britz test dataset possesses more samples than other test sites and also achieved the 0.5 threshold. This test dataset does however have the same trees in the training dataset giving the model an advantage to the other test sites.

**Figure 18:**
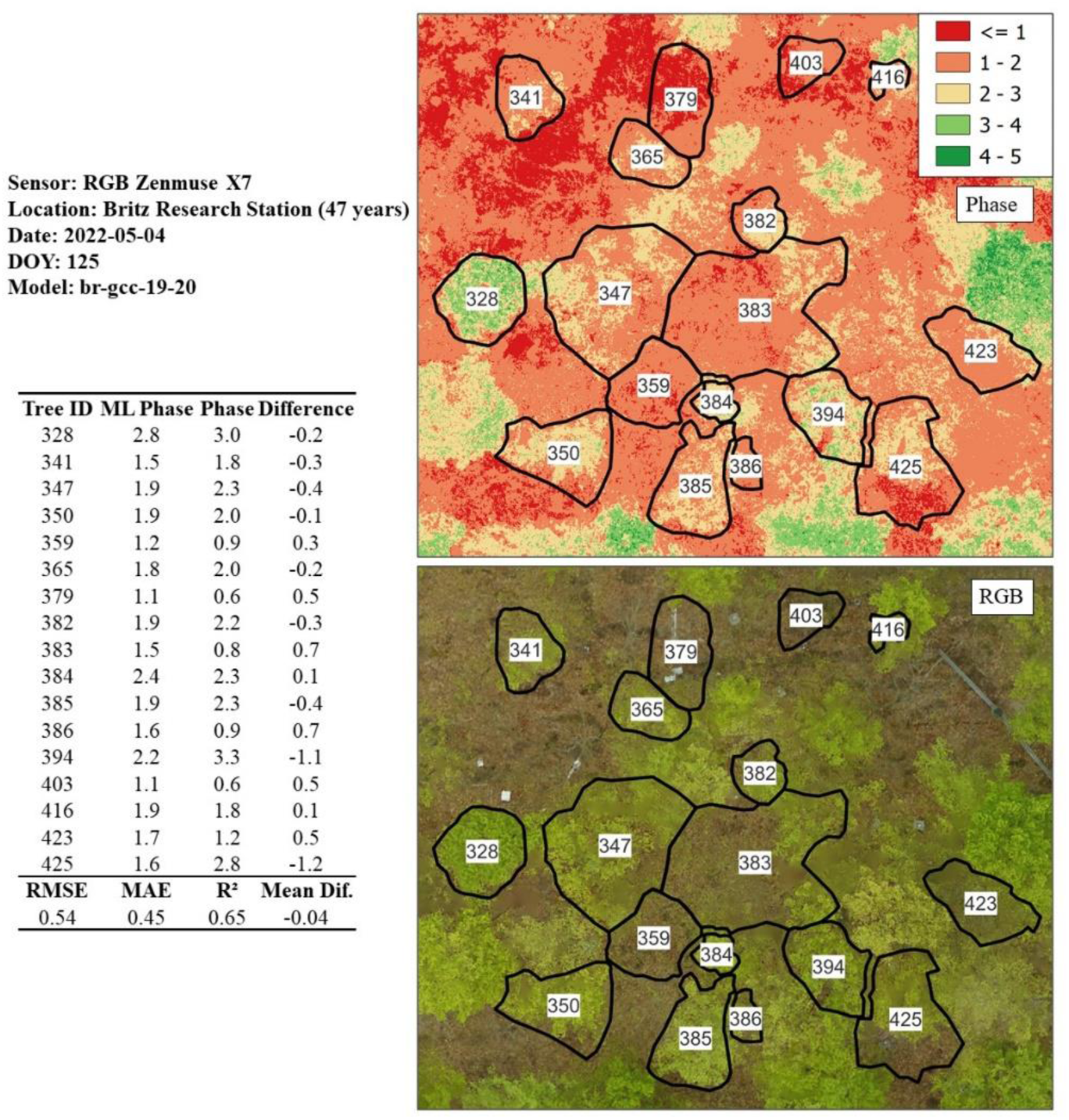
Phase prediction of a Beech stand (47 years) utilising the model originating from the calibrated GCC 2020/2021 dataset. Despite being a larger dataset (n = 17) in comparison to the other test sites, a RMSE of 0.54 was achieved which is can be regarded as achieving the 0.5 threshold.

**Figure 19:**
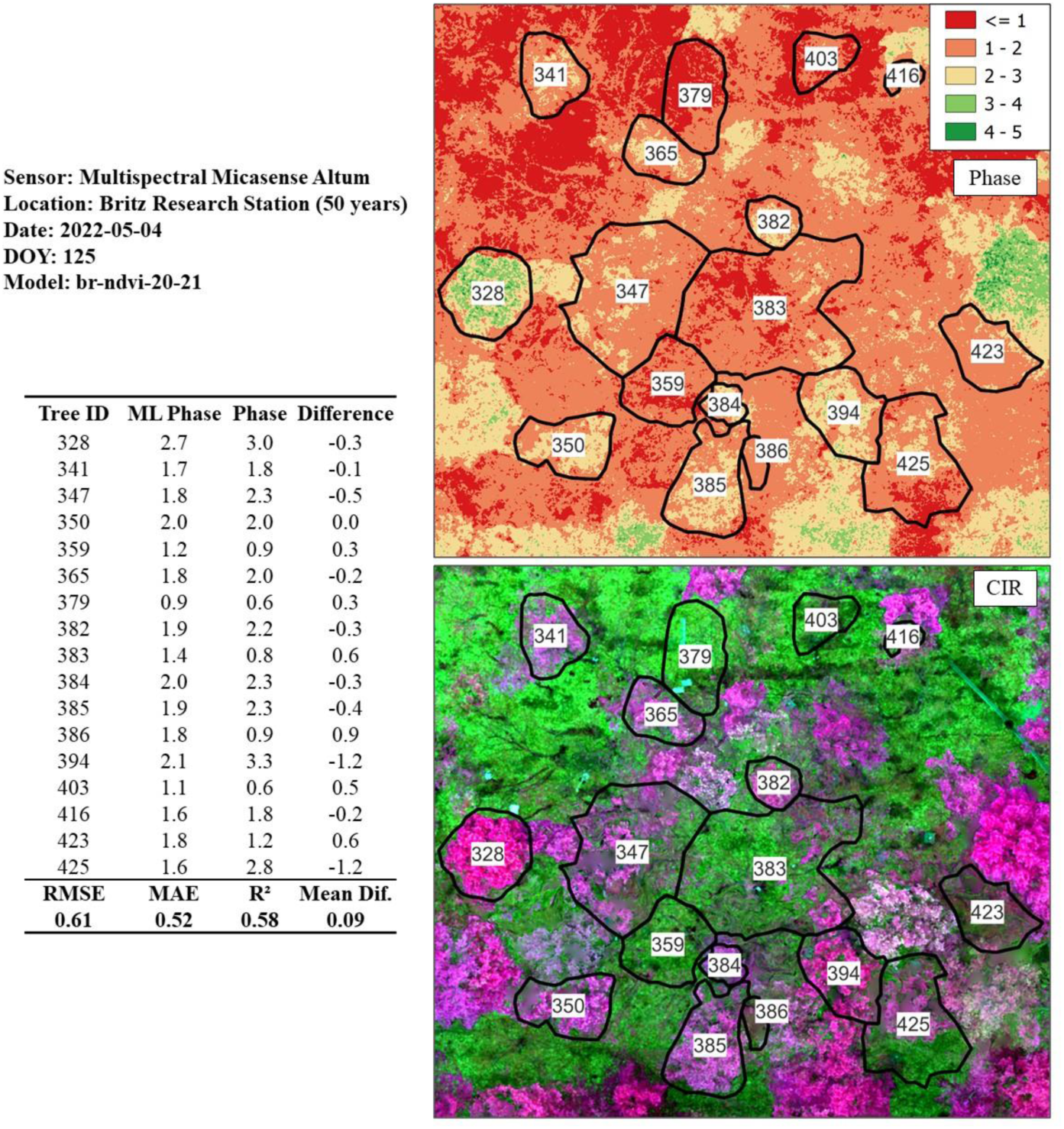
Phase prediction of a Beech stand (50 years) utilising the model originating from the calibrated NDVI 2020/2021 dataset. This is the only model derived from the non-visible band (NIR) which is in the proximity of the 0.5 threshold RMSE = 0.61). CIR = Color-infrared.

With respect to the test sites involving phase prediction from the multispectral sensor (*Micasesense Altum*), only the Britz and Kahlenberg sites were available. The only NDVI-based model that was in the proximity of the 0.5 threshold was Britz test dataset (*br-ndvi-20-21)* with a RMSE of 0.61, MAE of 0.52, and R-squared of 0.58 (*n* = 17). We hypothesized that the radiometric calibration methods from 2019 will have an effect on the model accuracy, however there was only a marginal difference in RMSE of the 2019/2020 and 2020/21 datasets.

Overall, the best performing and most consistent model for the purpose of predicting the spring phenological phases was that of the calibrated GCC based on the 2019/2020 training dataset. This model was capable of generalizing well over the test sites with the highest RMSE being from the Britz 2022 test site (RMSE = 0.54).

### 3.6 Sources of Error and Synopsis

This study brings to light the difficulties in acquiring radiometrically calibrated datasets of longer time series over multiple growing seasons despite before and after mission calibration panel acquisition and DLS data usage. The problem lies in the bottoming out of reflectance values, when for example the NDVI or any other indices involving the NIR band is calculated and occurs when during the flight mission the clouds open up only temporarily revealing the direct sun. Similar difficulties of oversaturation in the NIR band was also reported by Wang (2021). Sudden changes in illumination can influence several images or whole flight lines, and cannot be accounted for during calibration panel acquisition typically carried out according to the manufacturer (*micasense.com*) before and after the mission. The DLS does account for fluctuations in irradiance however only effectively for global changes in lighting conditions (*micasense.com*). This problem is further amplified in dense forests where the acquisition of shadow free reference panels is nearly impossible and capturing calibration data at a different location before and after missions is impractical and could result in an increase time differences to the actual flight mission where considerable changes in Solar angle could occur. The mitigation of oversaturation could potentially be accomplished with the manual experimentation of exposure settings however the challenge of acquiring reliable reflectance panel data still remains a challenge.

The factor involving the difficulty of radiometric calibration could lie also in the size of the reflectance panels themselves. Honkavaara et al., (2020) showed that larger custom-made reference panels of 1 x 1 meters calibrated better that the manufacturer’s provided method. Some studies also demonstrated improved calibration methods using even larger reflectance tarps (Honkavaara et al., 2012; H. Li et al., 2015; Moran et al., 2001) however still does not alleviate the problem of acquiring calibration data in dense forests as well as previously mentioned sudden changes illumination. Further testing, and the development of an improved field radiometric calibration strategy is an imperative in order to more efficiently exploit the capabilities of multispectral sensors.

Despite the poor results of the multispectral sensors, at least in the NIR band usage in this study, the utility of the RGB bands is somewhat significant. Typical low-cost UAV setups mounted with RGB sensors are widely available, which increases the possibility for attaining copious amounts of data. High data volume and availability will be an imperative when developing models for all of the relevant tree species available at intensive monitoring plots. The question here lies in if training data for models derived from visible bands require to be calibrated from the multispectral sensor? Here, the model trained with the calibrated GCC generalized well with the uncalibrated GCC, however will this be the case with new datasets as well as other tree species?

Another source of error could also arise in terms of the crown segmentation for the extraction of pixel values. Here for example a neighboring tree with an earlier phenological onset could have branches “bleeding” over into the segmented area of the crown in question. As manual or automatic segmentation is typically carried with a fully developed canopy (after phase 5.0), such overlapping branches will be difficult to account for. For the acquisition of high-quality training data, such influential branches from neighboring trees could be recorded during the ground observations and excluded from training datasets.

The feature selection process implemented in this study, in particular the partitioning of training datasets by year for further testing proved effective as it was possible to scrutinize and essentially remove portions of training data which could be having an effect on the generalisability of models. For example, the *br-ndvi-20-21* model derived from the multispectral sensors excludes the 2019 dataset which had lower quality radiometric calibration, time differences between observations, slightly different multispectral sensor and different observer for ground observations. On the other hand, the *gcc-19-20* models had generalized well with the 2019 datasets incorporated, however here only bands from the visible parts of the spectrum are used. We could deduce at this point that the main factors in error propagation lie in the quality of radiometric calibration and sensor mixing with the NIR bands which would not necessarily been apparent without the partitioning of training by year. Sensor mixing does however not seem to be an issue with RGB imagery which is advantageous for the acquisition large amounts of data.

With regards to the incorporation of meteorological data, as experimented with the usage of the “warming” days (AIRTEMP) as a model feature, we can deduce that other factors should also be considered here in terms of a dynamic start date and chilling days for a successful phenological model in fusion with spectral data. This concept is however still somewhat limited, as at the individual tree level, meteorological data will theoretically not be capable in explaining the heterogeneity of individual trees in terms of phenological development. The fusion of meteorological data alongside spectral data would however be reserved for larger scale applications, where phenological data is implemented stand-wise rather than on the individual tree level.

In terms of the *Britzer* foliation method, a translation of ground observations into remote sensing data was not possible due to the highly subject nature of this specific foliation estimation method. For this reason, among others, the *Britzer* method of foliation has been abandoned at the Brtiz research station and replaced with the ICP flushing method. Currently, the long-term *Britzer* phase method, alongside the flushing method are carried out with the aim of simplifying observations and enabling a harmonization of Britz research station data with the ICP network at the international level.

## 4 Conclusion and Future Outlook

Our research involves a machine learning approach for spring phenological phase prediction of European beech using UAV multispectral data. Three years (2019 – 2021) of synchronous ground observation and UAV-derived multisensor indices were applied to train and validate various machine learning models. A rigorous method of feature selection was introduced using Spearman correlation, polynomial fitting as well as machine learning. Further examination of models was carried out on unseen data while testing various training data partitions by year for the purpose of determining sources of error. The most successful training data partition, vegetation index and machine learning algorithm combination was that of the 2019/2020 dataset alongside the GCC and GAM boosting. The model generalized well with unseen data with a RMSE of 0.22 at the Kahlenberg site, 0.43 at the Black Forest site and 0.54 with data from 2022 at the Britz Research Station. The Britzer method of foliation was not able to be modelled successfully with RMSE values resulting well over the 10 % error threshold. Our findings highlight the development of a feature selection-based machine learning pipeline that utilizes radiometrically calibrated visible bands capable in predicting spring phenological phases with RGB imagery derived from readily available low-cost sensors.

The usage of the *Britzer* phenological phase method is of particular use at intensive monitoring plots such as the Britz research station, however the method is somewhat comprehensive for external plots. Here, the ICP Forests flushing method would be of utility as the incorporation of UAV-based data could enhance available datasets with phenology flushing (ICP Forests) predictions stand-wise and over larger areas. Large scale mapping of phenological flushing could in turn enable the training and validation of models based on data acquired from satellite platforms for the purpose of upscaling. This could facilitate not only large-scale mapping applications of forest phenology but also the creation of historical phenological time-series maps with the aim of assessing the effects of climate change with a spatial component. Nevertheless, extensive research and experimentation is still required and we recommend the following topics for future research:

- Focus on visible bands (RGB) as data is typically plentiful and test for the further feasibility of sensor mixing.
- Training data acquisition campaigns where an even distribution of particular phases is achieved. Here a well communicated and synchronous cooperation with ground observation crews is an imperative.
- Model development for further tree species available at intensive monitoring plots.
- Adaption of ICP Forests flushing levels to UAV-based modelling on an international level. Here regionalization of models will become a factor.
- Special training data acquisition campaigns for the training of flushing models involving ground observation experts could be applied for acquiring more continuous flushing values (i.e. 5 % levels) rather than relying on five code classes for the training of models. This would enable the usage of regression-based machine learning models which prove more effective than classification-based models and enable high quality training data.
- Development of models to predict other phase or flushing levels from a single epoch. This aspect is significant for acquiring more accurate timing of when flushing or particular phases occur rather than relying on statistical interpolation methods.
- Translation of observations or machine learning predictions to the stand-level for model development for Satellite applications.

## Acknowledgments

We would like to thank the team at the Britz Research Station (Thünen Institute of Forest Ecosystems) in particular Dietmar Fenske for assistance in the field as well as Prof. Dr. Jan-Peter Mund and the University for Sustainable development (HNEE) for support with equipment. Thanks also to Dr. Inken Krüger (Thünen-Institut) and Tilman Bucher (DLR) for reviewing the Manuscript.

## Data Availability

The UAV-based spring phenology model alongside sample workflow from this study can be found at https://git-dmz.thuenen.de/krause/uav_phenology and implemented for testing purposes.

### Declarations

### Author Contributions Statement

S.K. and T.S. designed the experiments. S.K. carried out the experiments, analysis and wrote the manuscript. T.S. contributed to the final version of the manuscript and supervised the project.

## References

Ahrends, H. E., Brügger, R., Stöckli, R., Schenk, J., Michna, P., Jeanneret, F., Wanner, H., & Eugster, W. (2008). Quantitative phenological observations of a mixed beech forest in northern Switzerland with digital photography. Journal of Geophysical Research: Biogeosciences, 113(G4). https://doi.org/10.1029/2007JG000650

Alberton, B., Torres, R. da S., Cancian, L. F., Borges, B. D., Almeida, J., Mariano, G. C., Santos, J. dos, & Morellato, L. P. C. (2017). Introducing digital cameras to monitor plant phenology in the tropics: Applications for conservation. Perspectives in Ecology and Conservation, 15(2), 82–90. https://doi.org/10.1016/j.pecon.2017.06.004

Atkins, J. W., Stovall, A. E. L., & Yang, X. (2020). Mapping Temperate Forest Phenology Using Tower, UAV, and Ground-Based Sensors. Drones, 4(3), 56. https://doi.org/10.3390/drones4030056

Augspurger, C. K. (2009). Spring 2007 warmth and frost: Phenology, damage and refoliation in a temperate deciduous forest. Functional Ecology, 23(6), 1031–1039. https://doi.org/10.1111/j.1365-2435.2009.01587.x

Awaya, Y., Tanaka, K., Kodani, E., & Nishizono, T. (2009). Responses of a beech (Fagus crenata Blume) stand to late spring frost damage in Morioka, Japan. Forest Ecology and Management, 257(12), 2359–2369. https://doi.org/10.1016/j.foreco.2009.03.028

Barnes, E. M., Clarke, T. R., Richards, S. E., Colaizzi, P. D., Haberland, J., Kostrzewski, M., Waller, P., Choi, C., Riley, E., Thompson, T., Lascano, R. J., Li, H., & Moran, M. S. (2000). COINCIDENT DETECTION OF CROP WATER STRESS, NITROGEN STATUS AND CANOPY DENSITY USING GROUND-BASED MULTISPECTRAL DATA. 16.

Baumgartner, A. (1952). Zur Phänologie von Laubhölzern und ihre Anwendung bei lokalklimatischen Untersuchungen. Berichte Des DWD in Der US-Zone, 42, 69–73.

Belle, V., & Papantonis, I. (2021). Principles and Practice of Explainable Machine Learning. Frontiers in Big Data, 4. https://www.frontiersin.org/articles/10.3389/fdata.2021.688969

Berra, E. F., Gaulton, R., & Barr, S. (2019). Assessing spring phenology of a temperate woodland: A multiscale comparison of ground, unmanned aerial vehicle and Landsat satellite observations. Remote Sensing of Environment, 223, 229–242. https://doi.org/10.1016/j.rse.2019.01.010

Brown, T. B., Hultine, K. R., Steltzer, H., Denny, E. G., Denslow, M. W., Granados, J., Henderson, S., Moore, D., Nagai, S., SanClements, M., Sánchez-Azofeifa, A., Sonnentag, O., Tazik, D., & Richardson, A. D. (2016). Using phenocams to monitor our changing Earth: Toward a global phenocam network. Frontiers in Ecology and the Environment, 14(2), 84–93. https://doi.org/10.1002/fee.1222

Brügger, R., & Vasella, A. (2018). Pflanzen im Wandel der Jahreszeiten. Anleitung für phänologische Beobachtungen / Les plantes au cours des saisons. Guide pour observation phénologiques. Geographica Bernensia. http://doi.org/10.4480/GB2018.N02

Chandrashekar, G., & Sahin, F. (2014). A survey on feature selection methods. Computers & Electrical Engineering, 40(1), 16–28. https://doi.org/10.1016/j.compeleceng.2013.11.024

Cleland, E., Chuine, I., Menzel, A., Mooney, H., & Schwartz, M. (2007). Shifting plant phenology in response to global change. Trends in Ecology & Evolution, 22(7), 357–365. https://doi.org/10.1016/j.tree.2007.04.003

Copernicus. (2022). Sentinel Online—ESA - Sentinel Online. https://sentinels.copernicus.eu/web/sentinel/home

Czernecki, B., Nowosad, J., & Jabłońska, K. (2018). Machine learning modeling of plant phenology based on coupling satellite and gridded meteorological dataset. International Journal of Biometeorology, 62(7), 1297–1309. https://doi.org/10.1007/s00484-018-1534-2

Dai, W., Jin, H., Zhang, Y., Liu, T., & Zhou, Z. (2019). Detecting temporal changes in the temperature sensitivity of spring phenology with global warming: Application of machine learning in phenological model. Agricultural and Forest Meteorology, 279, 107702. https://doi.org/10.1016/j.agrformet.2019.107702

Dethier, B. E., Ashley, M. D., & Blair, B. (1972). PHENOLOGY SATELLITE EXPERIMENT. 9.

Diez, Y., Kentsch, S., Fukuda, M., López Caceres, M. L., Moritake, K., & Cabezas, M. (2021). Deep Learning in Forestry Using UAV-Acquired RGB Data: A Practical Review. Remote Sensing, 13, 2837. https://doi.org/10.3390/rs13142837

Don, A., Hagen, C., Grüneberg, E., & Vos, C. (2019). Simulated wild boar bioturbation increases the stability of forest soil carbon. Biogeosciences, 16(21), 4145–4155. https://doi.org/10.5194/bg-16-4145-2019

Forstreuter, M. (2002). Auswirkungen globaler Klimaänderungen auf das Wachstum und den Gaswechsel (CO2/H2O) von Rotbuchenbeständen (Fagus sylvatica L.). Techn. Univ.

Friedl, M., Sulla-Menashe, D., Tan, B., Schneider, A., Ramankutty, N., Sibley, A., & Huang, X. (2010). MODIS Collection 5 Global Land Cover: Algorithm Refinements and Characterization of new Datasets. Remote Sensing of Environment, 114, 168–182. https://doi.org/10.1016/j.rse.2009.08.016

Ganguly, S., Friedl, M. A., Tan, B., Zhang, X., & Verma, M. (2010). Land surface phenology from MODIS: Characterization of the Collection 5 global land cover dynamics product. Remote Sensing of Environment, 114(8), 1805–1816. https://doi.org/10.1016/j.rse.2010.04.005

Gillespie, A. R., Kahle, A. B., & Walker, R. E. (1987). Color enhancement of highly correlated images. II. Channel ratio and “chromaticity” transformation techniques. Remote Sensing of Environment, 22(3), 343–365. https://doi.org/10.1016/0034-4257(87)90088-5

Gitelson, A. A., Kaufman, Y. J., & Merzlyak, M. N. (1996). Use of a green channel in remote sensing of global vegetation from EOS-MODIS. Remote Sensing of Environment, 58(3), 289–298. https://doi.org/10.1016/S0034-4257(96)00072-7

Gitelson, A., & Merzlyak, M. (1994a). Quantitative estimation of chlorophyll-a using reflectance spectra: Experiments with autumn chestnut and maple leaves. Journal of Photochemistry and Photobiology B: Biology, 22, 247–252. https://doi.org/10.1016/1011-1344(93)06963-4

Gitelson, A., & Merzlyak, M. N. (1994b). Quantitative estimation of chlorophyll-a using reflectance spectra: Experiments with autumn chestnut and maple leaves. Journal of Photochemistry and Photobiology B: Biology, 22(3), 247–252. https://doi.org/10.1016/1011-1344(93)06963-4

Honkavaara, E., Hakala, T., Markelin, L., Rosnell, T., Saari, H., & Mäkynen, J. (2012). A Process for Radiometric Correction of UAV Image Blocks. Photogrammetrie - Fernerkundung - Geoinformation, 2012(2), 115–127. https://doi.org/10.1127/1432-8364/2012/0106

Honkavaara, E., Näsi, R., Alves de Oliveira, R., Viljanen, N., Suomalainen, J., Khoramshahi, E., Hakala, T., Nevalainen, O., Markelin, L., Vuorinen, M., Kankaanhuhta, V., Paivi, L.-S., & Haataja, L. (2020). Using Multitemperaol Hyper- and Multispectral UAV Imaging for Detecting Bark Beetle Infestation on Norway Spruce. ISPRS - International Archives of the Photogrammetry, Remote Sensing and Spatial Information Sciences, XLIII-B3-2020, 429–434. https://doi.org/10.5194/isprs-archives-XLIII-B3-2020-429-2020

Huete, A., Didan, K., Miura, T., Rodriguez, E. P., Gao, X., & Ferreira, L. G. (2002). Overview of the radiometric and biophysical performance of the MODIS vegetation indices. Remote Sensing of Environment, 83(1–2), 195–213.

Hunt, E. R., Doraiswamy, P. C., McMurtrey, J. E., Daughtry, C. S. T., Perry, E. M., & Akhmedov, B. (2013). A visible band index for remote sensing leaf chlorophyll content at the canopy scale. International Journal of Applied Earth Observation and Geoinformation, 21, 103– 112. https://doi.org/10.1016/j.jag.2012.07.020

IPCC. (2018). Technical Summary—Special Report on Climate Change and Land. https://www.ipcc.ch/srccl/chapter/technical-summary/

Jones, H. G., & Vaughan, R. A. (2010). Remote Sensing of Vegetation: Principles, Techniques, and Applications. Oxford University Press.

Kattenborn, T., Leitloff, J., Schiefer, F., & Hinz, S. (2021). Review on Convolutional Neural Networks (CNN) in vegetation remote sensing. ISPRS Journal of Photogrammetry and Remote Sensing, 173, 24–49. https://doi.org/10.1016/j.isprsjprs.2020.12.010

Klosterman, S., Melaas, E., Wang, J. A., Martinez, A., Frederick, S., O’Keefe, J., Orwig, D. A., Wang, Z., Sun, Q., Schaaf, C., Friedl, M., & Richardson, A. D. (2018). Fine-scale perspectives on landscape phenology from unmanned aerial vehicle (UAV) photography. Agricultural and Forest Meteorology, 248, 397–407. https://doi.org/10.1016/j.agrformet.2017.10.015

Klosterman, S. T., Hufkens, K., Gray, J. M., Melaas, E., Sonnentag, O., Lavine, I., Mitchell, L., Norman, R., Friedl, M. A., & Richardson, A. D. (2014). Evaluating remote sensing of deciduous forest phenology at multiple spatial scales using PhenoCam imagery. Biogeosciences, 11(16), 4305–4320. https://doi.org/10.5194/bg-11-4305-2014

Kowalski, K., Senf, C., Hostert, P., & Pflugmacher, D. (2020). Characterizing spring phenology of temperate broadleaf forests using Landsat and Sentinel-2 time series. International Journal of Applied Earth Observation and Geoinformation, 92, 102172. https://doi.org/10.1016/j.jag.2020.102172

Kuhn, M. (2008). Building Predictive Models in R Using the caret Package. Journal of Statistical Software, 28(5). https://doi.org/10.18637/jss.v028.i05

Kuhn, M., & Johnson, K. (2019). Feature Engineering and Selection: A Practical Approach for Predictive Models. CRC Press.

Kuhn, M., Wing, J., Weston, S., & Williams, A. (2022). The caret package. Gene Expr. Landsat. (2022). Landsat Science. https://landsat.gsfc.nasa.gov/

Lary, D. J., Alavi, A. H., Gandomi, A. H., & Walker, A. L. (2016). Machine learning in geosciences and remote sensing. Geoscience Frontiers, 7(1), 3–10. https://doi.org/10.1016/j.gsf.2015.07.003

Li, H., Zhang, H., Chen, Z., Yang, M., & Zhang, Y. (2015). A Method Suitable for Vicarious Calibration of a UAV Hyperspectral Remote Sensor. IEEE Journal of Selected Topics in Applied Earth Observations and Remote Sensing, 8, 1–15. https://doi.org/10.1109/JSTARS.2015.2416213

Li, N., Zhan, P., Pan, Y., Zhu, X., Li, M., & Zhang, D. (2020). Comparison of Remote Sensing Time-Series Smoothing Methods for Grassland Spring Phenology Extraction on the Qinghai–Tibetan Plateau. Remote Sensing, 12(20), 3383. https://doi.org/10.3390/rs12203383

Liang, S., & Wang, J. (2020). Advanced remote sensing: Terrestrial information extraction and applications.

Lieth, H. (Ed.). (1974). Phenology and Seasonality Modeling (Vol. 8). Springer Berlin Heidelberg. https://doi.org/10.1007/978-3-642-51863-8

Linderholm, H. W. (2006). Growing season changes in the last century. Agricultural and Forest Meteorology, 137(1–2), 1–14. https://doi.org/10.1016/j.agrformet.2006.03.006

Linnaeus, C. (1751). Philosophia botanica: In qua explicantur fundamenta botanica cum definitionibus partium, exemplis terminorum, observationibus rariorum, adjectis figuris aeneis. apud Godofr. Kiesewetter.

Malaisse, F. (1964). CONTRIBUTION A L’ÉTUDE DES HÊTRAIES D’EUROPE OCCIDENTALE: Note 4: Quelques observations phénologiques de hêtraies en 1963. Bulletin de La Société Royale de Botanique de Belgique / Bulletin van de Koninklijke Belgische Botanische Vereniging, 97, 85–97. JSTOR.

Mather, P. M., & Koch, M. (2011). Computer processing of remotely-sensed images: An introduction (4th ed). Wiley-Blackwell.

Maxwell, A. E., Warner, T. A., & Fang, F. (2018). Implementation of machine-learning classification in remote sensing: An applied review. International Journal of Remote Sensing, 39(9), 2784–2817. https://doi.org/10.1080/01431161.2018.1433343

McClave, J. T., & Sincich, T. T. (2018). Statistics, Global Edition (13. Edition). Pearson Education Limited.

McFeeters, S. K. (1996). The use of the Normalized Difference Water Index (NDWI) in the delineation of open water features. International Journal of Remote Sensing, 17(7), 1425– 1432. https://doi.org/10.1080/01431169608948714

Menzel, A. (1997). Phänologie von Waldbäumen unter sich ändernden Klimabedingungen: Auswertung der Beobachtungen in den internationalen phänologischen Gärten und Möglichkeiten der Modellierung von Phänodaten. Frank.

Menzel, A. (2002). Phenology: Its Importance to the Global Change Community. Climatic Change, 54(4), 379–385. https://doi.org/10.1023/A:1016125215496

Menzel, A., Helm, R., & Zang, C. (2015). Patterns of late spring frost leaf damage and recovery in a European beech (Fagus sylvatica L.) stand in south-eastern Germany based on repeated digital photographs. Frontiers in Plant Science, 6. https://doi.org/10.3389/fpls.2015.00110

Menzel, A., Sparks, T. H., Estrella, N., Koch, E., Aasa, A., Ahas, R., Alm-Kübler, K., Bissolli, P., Braslavská, O., Briede, A., Chmielewski, F. M., Crepinsek, Z., Curnel, Y., Dahl, Å., Defila, C., Donnelly, A., Filella, Y., Jatczak, K., Måge, F., … Zust, A. (2006). European phenological response to climate change matches the warming pattern. Global Change Biology, 12(10), 1969–1976. https://doi.org/10.1111/j.1365-2486.2006.01193.x

micasense.com. (2022). *Micasense.com*. https://micasense.com/

Moran, M. S., Bryant, R. B., Clarke, T. R., & Qi, J. (2001). Deployment and calibration of reference reflectance tarps for use with airborne imaging sensors. Photogrammetric Engineering and Remote Sensing, 67(3), 273–286.

MüllerA, J. (2010). Forest hydrology research with lysimeter in the northeast German lowlands special methods and results for the forest management. Soil Solutions for a Changing World: Proceedings of the 19th World Congress of Soil Science, Edited by RJ Gilkes and N. Prakongkep, 28–31. https://www.researchgate.net/profile/Andreas_Bolte/publication/265980128_Forest_hydrology_research_with_lysimeter_in_the_northeast_German_lowlands_special_methods_and_results_for_the_forest_management/links/54d2097d0cf28370d0e199cb.pdf

Park, J. Y., Muller-Landau, H. C., Lichstein, J. W., Rifai, S. W., Dandois, J. P., & Bohlman, S. A. (2019). Quantifying Leaf Phenology of Individual Trees and Species in a Tropical Forest Using Unmanned Aerial Vehicle (UAV) Images. Remote Sensing, 11(13), 1534. https://doi.org/10.3390/rs11131534

R Core Team. (2022). R: A language and environment for statistical computing. R Foundation for Statistical Computing, Vienna, Austria. http://www.R-project.org/

Raspe, S., Fleck, S., Beuker, E., Bastrup-Birk, A., & Preuhsler, T. (2020). Manual on methods and criteria for harmonized sampling, assessment, monitoring and analysis of the effects of air pollution on forests. Thünen Institute of Forest Ecosystems, Eberswalde, Germany. http://www.icp-forests.org/Manual.htm

Riek, W. (2004). Eigenschaften typischer Waldböden im Nordostdeutschen Tiefland unter besonderer Berücksichtigung des Landes Brandenburg: Vol. Bd. 19. Ministerium für Landwirtschaft, Umweltschutz und Raumordnung des Landes Brandenburg, Presse- und Öffentlichkeitsarbeit [u.a.]. http://www.worldcat.org/oclc/64657313

Rodrigues, A., Marcal, A. R. S., & Cunha, M. (2012). Phenology parameter extraction from time-series of satellite vegetation index data using phenosat. 2012 IEEE International Geoscience and Remote Sensing Symposium, 4926–4929. https://doi.org/10.1109/IGARSS.2012.6352507

Rouse, J. W., Haas, R. H., Schell, J. A., Deering, D. W., & Harlan, J. C. (1974). Monitoring the vernal advancements and retrogradation. Texas, Texas A & M University.

Rubio-Cuadrado, Á., Camarero, J. J., Rodríguez-Calcerrada, J., Perea, R., Gómez, C., Montes, F., & Gil, L. (2021). Impact of successive spring frosts on leaf phenology and radial growth in three deciduous tree species with contrasting climate requirements in central Spain. Tree Physiology, 41(12), 2279–2292. https://doi.org/10.1093/treephys/tpab076

Sachsenforst. (2020, June 25). Frost trifft Forst. https://medienservice.sachsen.de/medien/news/238026

Sakai, A., & Larcher, W. (1987). Frost Survival of Plants (Vol. 62). Springer Berlin Heidelberg. https://doi.org/10.1007/978-3-642-71745-1

Schuldt, B., Buras, A., Arend, M., Vitasse, Y., Beierkuhnlein, C., Damm, A., Gharun, M., Grams, T. E. E., Hauck, M., Hajek, P., Hartmann, H., Hiltbrunner, E., Hoch, G., Holloway-Phillips, M., Körner, C., Larysch, E., Lübbe, T., Nelson, D. B., Rammig, A., … Kahmen, A. (2020). A first assessment of the impact of the extreme 2018 summer drought on Central European forests. Basic and Applied Ecology, 45, 86–103. https://doi.org/10.1016/j.baae.2020.04.003

Schüler, S. (2012). Genetische Variation und Plastizität des Blattaustriebs von Herkünften der Rot-Buche. 10.

Schulz, K., Hänsch, R., & Sörgel, U. (2018). Machine learning methods for remote sensing applications: An overview. Earth Resources and Environmental Remote Sensing/GIS Applications IX, 10790, 1079002. https://doi.org/10.1117/12.2503653

Schulze, G., & Kopp, D. (1998). Anleitung für die forstliche Standortserkundung im nordostdeutschen Tiefland (Standortserkundungsanleitung) SEA 95, Teil C–Forstliche Auswertung. Bodenformen-Katalog. Merkmalsübersichten Und-Tabellen Für Haupt-Und Feinbodenformen. Unter Mitarbeit von D. Kopp, 3.

Schwartz, M. D. (Ed.). (2013). Phenology: An Integrative Environmental Science. Springer Netherlands. https://doi.org/10.1007/978-94-007-6925-0

Sonnentag, O., Hufkens, K., Teshera-Sterne, C., Young, A. M., Friedl, M., Braswell, B. H., Milliman, T., O’Keefe, J., & Richardson, A. D. (2012). Digital repeat photography for phenological research in forest ecosystems. Agricultural and Forest Meteorology, 152, 159–177. https://doi.org/10.1016/j.agrformet.2011.09.009

Tucker, C. J. (1979). Red and photographic infrared linear combinations for monitoring vegetation. Remote Sensing of Environment, 8(2), 127–150.

Vilhar, U., Beuker, E., Mizunuma, T., Skudnik, M., Lebourgeois, F., Soudani, K., & Wilkinson, M. (2013). Tree Phenology. In Developments in Environmental Science (Vol. 12, pp. 169–182). Elsevier. https://doi.org/10.1016/B978-0-08-098222-9.00009-1

Wang, C. (2021). At-Sensor Radiometric Correction of a Multispectral Camera (RedEdge) for sUAS Vegetation Mapping. Sensors, 21(24), Article 24. https://doi.org/10.3390/s21248224

White, K., Pontius, J., & Schaberg, P. (2014). Remote sensing of spring phenology in northeastern forests: A comparison of methods, field metrics and sources of uncertainty. Remote Sensing of Environment, 148, 97–107. https://doi.org/10.1016/j.rse.2014.03.017

White, M. A., Running, S. W., & Thornton, P. E. (1999). The impact of growing-season length variability on carbon assimilation and evapotranspiration over 88 years in the eastern US deciduous forest. International Journal of Biometeorology, 42(3), 139–145. https://doi.org/10.1007/s004840050097

Zeng, L., Wardlow, B. D., Xiang, D., Hu, S., & Li, D. (2020). A review of vegetation phenological metrics extraction using time-series, multispectral satellite data. Remote Sensing of Environment, 237, 111511. https://doi.org/10.1016/j.rse.2019.111511

Zhang, X. (2012). Phenology and Climate Change. https://doi.org/10.5772/2146

Zhao, M., Peng, C., Xiang, W., Deng, X., Tian, D., Zhou, X., Yu, G., he, H., & Zhao, Z. (2013). Plant phenological modeling and its application in global climate change research: Overview and future challenges. Environmental Reviews, 21. https://doi.org/10.1139/er-2012-0036

Zhu, W., Mou, M., Wang, L., & Jiang, N. (2012). Evaluation of phenology extracting methods from vegetation index time series. 2012 IEEE International Geoscience and Remote Sensing Symposium, 1158–1161. https://doi.org/10.1109/IGARSS.2012.6351342

